# Myosin Heavy Chain-embryonic is a crucial regulator of skeletal muscle development and differentiation

**DOI:** 10.1101/261685

**Authors:** Akashi Sharma, Megha Agarwal, Amit Kumar, Pankaj Kumar, Masum Saini, Gabrielle Kardon, Sam J. Mathew

## Abstract

Myosin heavy chains (MyHCs) are contractile proteins that are part of the thick filaments of the functional unit of the skeletal muscle, the sarcomere. In addition to MyHCs that are part of the adult muscle contractile network, two MyHCs - MyHC-embryonic and -perinatal are expressed during muscle development and are only transiently expressed in the adult during regeneration. The functions performed by these MyHCs has been a long-standing question and using a targeted mouse allele, we have characterized the role of MyHC-embryonic. Analysis of loss-of-function mice reveals that lack of MyHC-embryonic leads to mis-regulation of other MyHCs, alterations in fiber size, fiber number and fiber type at neonatal stages. We also find that loss of MyHC-embryonic leads to mis-regulation of genes involved in muscle differentiation. A broad theme from these studies is that loss of MyHC-embryonic has distinct effects on different muscles, possibly reflecting the unique fiber type composition of different muscles. Most significantly, our results indicate that MyHC-embryonic is required during embryonic and fetal myogenesis to regulate myogenic progenitor and myoblast differentiation in a non-cell autonomous manner via Mitogen Activated Protein Kinase (MAPKinase) and Fibroblast Growth Factor (FGF) signaling. Thus, our results signify that MyHC-embryonic is a key regulator of myogenic differentiation during embryonic, fetal and neonatal myogenesis.

## Introduction

The vertebrate skeletal muscle is a contractile tissue that develops through a step-wise process of differentiation from progenitor cells during development. During vertebrate development, the skeletal muscle arises from the mesoderm, with the trunk and limb musculature derived from the dorsal compartment of the somites, the dermomyotome (Braun and Gautel, 2011; Buckingham, 1992). The myogenic progenitors (MPs) formed in the somites migrate into target organs such as the limbs and the diaphragm (Dietrich, et al., 1999; Bladt, et al., 1995). MPs undergo a differentiation program, thought to occur in two phases during mouse development-an embryonic phase and a fetal phase (Biressi, et al., 2007a; Biressi, et al., 2007b; Stockdale, 1992). During the embryonic phase of myogenesis (Embryonic day [E] 9.5-13.5 in mice), a small proportion of the embryonic MPs fuse with each other to give rise to primary myofibers, while in the fetal phase (E14.5-17.5), fetal MPs fuse with each other and with primary myofibers to give rise to secondary myofibers (Murphy and Kardon, 2011; Zammit, et al., 2008; Biressi, et al., 2007a; Biressi, et al., 2007b; Kelly, et al., 1997). While embryonic and fetal MPs differ from each other with respect to their morphology, proliferation rates, and response to growth factors, the myofibers derived from them, namely primary and secondary myofibers respectively, also exhibit distinct characteristics (Zammit, et al., 2008; Biressi, et al., 2007a; Pin, et al., 2002; Stockdale, 1992; Barbieri, et al., 1990; Cossu and Molinaro, 1987). Primary myofibers are fewer in number, while secondary myofibers are smaller and form around primary fibers (Biressi, et al., 2007a; Cossu and Biressi, 2005). Following the embryonic and fetal myogenic phases, perinatal and adult myogenesis takes place, during which growth, maturation, and functional diversification of the myofibers occurs. In adult mice, the MPs are called satellite cells residing between the myofiber plasma membrane and the basal lamina. Satellite cells are normally quiescent, but give rise to myofibers during muscle regeneration and contribute to myofibers during homeostasis (Keefe, et al., 2015; Murphy and Kardon, 2011; Biressi, et al., 2007a).

The transcriptional network involved in myogenesis has been well studied (Braun and Gautel, 2011; Murphy and Kardon, 2011). The MPs express two paired domain homeobox transcription factors, Pax3 and Pax7, which are critical for myogenesis (Braun and Gautel, 2011; Bryson-Richardson and Currie, 2008; Buckingham, 2007). In the limb, Pax3^+^ Pax7^−^ cells are required for embryonic myogenesis and to give rise to Pax7^+^ cells, while Pax3-derived, Pax7^+^ cells are required for fetal myogenesis, suggesting that these transcription factors play distinct roles in embryonic and fetal myogenesis (Hutcheson, et al., 2009). Downstream of Pax3 and Pax7 are four basic helix-loop-helix transcription factors termed myogenic regulatory factors (MRFs), which facilitate myogenic determination and terminal differentiation (Maroto, et al., 1997). They are Myf5, MyoD, MRF4 and Myogenin, with Myf5, and MyoD thought to be essential for myogenic determination, Myogenin required for terminal differentiation and MRF4 thought to play a dual determination-differentiation role (Braun and Gautel, 2011; Murphy and Kardon, 2011). MRFs, together with the Myocyte Enhancer Factor 2 (MEF2) family of transcription factors activate muscle specific target genes, which are required for sarcomeric organization and contractility (Bryson-Richardson and Currie, 2008).

The contractile proteins essential for muscle contraction include actin, myosin and associated proteins such as troponin, and tropomyosin (Gunning and Hardeman, 1991). The contractile unit of the skeletal muscle is the sarcomere, made up of primarily the thin and thick filaments composed of actin and myosin contractile proteins respectively, in addition to associated regulatory proteins (Braun and Gautel, 2011; Ehler and Gautel, 2008). The Myosin molecule is a heterohexamer comprising a pair of Myosin Heavy Chains (MyHCs), a pair of regulatory light chains and a pair of essential light chains. The two MyHCs have a N-terminal globular head region with actin-binding and ATPase activity, a neck region where the light chains bind and a C-terminal coiled-coil tail region (Schiaffino and Reggiani, 1996). Interestingly, it has been known that the contractile velocity of muscles is directly correlated with its myosin ATPase activity, and varies from muscle to muscle (Barany, 1967). Based on their contractile properties, MyHCs are broadly classified as fast or slow. There are mainly seven MyHC isoforms thought to be expressed in the mammalian skeletal muscle, of which five are expressed during adult life: MyHC-IIa, -IId and -IIb are adult “fast” isoforms, and MyHC-slow is the adult “slow” isoform; MyHC-extraocular is a unique “fast” isoform expressed in the extraocular and laryngeal muscles (Parker-Thornburg, et al., 1992; Narusawa, et al., 1987; Wieczorek, et al., 1985; Weydert, et al., 1983). Among these, MyHC-IIb expressing fibers exhibit fastest contractility, followed by MyHC-IIx, MyHC-IIa and MyHC-slow; MyHC-IIb and -IIx fibers majorly utilize glycolytic metabolism while MyHC-IIa and -slow fibers utilize oxidative metabolism. Thus, adult muscle fibers exhibit varying contractile properties, depending on which specific MyHCs are expressed in those fibers, the anatomical location and function of the muscle the fibers are a part of. Adult muscle fiber type has been extensively studied and numerous signals and pathways regulating adult muscle fiber type has been described (Schiaffino and Reggiani, 2011).

In addition to the five MyHC isoforms described above, two developmental isoforms MyHC-embryonic and -perinatal (MyHC-emb and -peri), are expressed during embryonic, fetal and neonatal development (Periasamy, et al., 1985; Periasamy, et al., 1984). Although MyHC-slow is expressed in the adult muscle, it is also expressed during developmental stages in the differentiated myofibers. MyHC-emb and -peri are not expressed in the mature adult muscle normally, except during skeletal muscle injury or disease where these two MyHCs are transiently re-expressed during regeneration (d’Albis, et al., 1988; Schiaffino, et al., 1986; Sartore, et al., 1982).

The precise mechanisms underlying fiber type specification during development has not been well characterized. The Six family of homeobox proteins have been reported to regulate the fast-type fiber specification during development (Niro, et al., 2010), while Sox6 is also thought to repress the slow-type fiber specification and activate fast fiber program during development (Hagiwara, et al., 2007; Hagiwara, et al., 2005). One critical factor required for the embryonic to fetal myogenic transition is the transcription factor Nfix, which is required to promote the fetal program, and repress MyHC-slow (Messina, et al., 2010). We had previously shown that extrinsic signals from the connective tissue fibroblasts are crucial for slow fiber specification during development, and fiber maturation (Mathew, et al., 2011). Thus, it is clear that both intrinsic and extrinsic signals are required for proper fiber type specification during development (Murphy and Kardon, 2011; Biressi, et al., 2007a). All muscle fibers express MyHC-slow during embryonic myogenesis, while during fetal myogenesis MyHC-slow is downregulated in most muscles and MyHC-perinatal is expressed; thus, embryonic myofibers are thought to be predominantly slow while fetal myofibers are predominantly fast (Schiaffino and Reggiani, 2011; Biressi, et al., 2007a). Each anatomical muscle exhibits a distinctive fiber type pattern during development and at birth, suggesting that there are tightly regulated mechanisms that regulate fiber type specification during development (Biressi, et al., 2007a). It is unknown whether MyHCs themselves are critical for determining fiber type during development.

Previously, studies on knockout mice for MyHC-IIb and IIx were carried out to understand the role of MyHCs in adult muscle function (Sartorius, et al., 1998; Acakpo-Satchivi, et al., 1997). Mice null for MyHC-IIb or -IIx were significantly smaller, weighed less than control animals, exhibiting reduced grip strength, muscle weakness and increased interstitial fibrosis (Acakpo-Satchivi, et al., 1997). While MyHC-IIb null mice had significantly reduced levels of muscle contractile force, MyHC-IIx null mice had altered kinetics of muscle contraction (Acakpo-Satchivi, et al., 1997). 6 week old MyHC-IIx null animals also exhibited kyphosis or curvature of the spinal column (Sartorius, et al., 1998). There are no knockout studies so far on any of the developmental MyHCs. Interestingly, mutations in the MyHC-emb encoding *MyH3* gene have been reported to cause Freeman-Sheldon and Sheldon-Hall congenital contracture syndromes while mutations in MyHC-peri encoding *MyH8* gene led to Trismus-pseudocamptodactyly congenital contracture syndrome in humans, indicating that developmental MyHCs have vital roles during embryonic development (Toydemir, et al., 2006b; Toydemir, et al., 2006c).

In this study, we characterize the role of MyHC-embryonic (MyHC-emb) in myogenic differentiation using a conditional targeted mouse allele during development in vivo, and C2C12 myogenic cells in vitro. We find that MyHC-emb protein is expressed in two peaks during mouse embryonic development and is expressed at considerably higher levels compared to MyHC-slow. Using a null allele and a conditional targeted allele for MyHC-emb, we investigate the role of MyHC-emb during embryonic, fetal and neonatal myogenesis. We find that MyHC-emb is a crucial developmental cue, required to regulate levels of other MyHCs, fiber type, fiber number and fiber area. Significantly, we find that although MyHC-emb is expressed in all myofibers during development, different muscles respond differently to the loss of MyHC-emb. We also find that in addition to cell autonomous roles in regulating myogenesis, MyHC-emb performs a novel, non-cell autonomous role in regulating the rate of myogenic progenitor and myoblast differentiation during development. Thus, our study shows for the first time that developmental MyHCs are important factors that play crucial cell autonomous and non-cell autonomous roles during animal development. Surprisingly, MyHC-emb expression in myofiber, controls a secreted signal, which feeds back and regulates myogenesis.

## Results

### MyHC-embryonic is expressed at high levels relative to MyHC-slow during embryonic, fetal and in vitro myogenesis

To characterize the expression of Myosin Heavy Chains (MyHCs) during mouse embryonic development, we quantified the transcript levels of MyHC-embryonic (MyHC-emb) and MyHC-slow starting at E 9.5, until P (postnatal day) 0, by qPCR. MyHC-emb transcript levels increase sharply from E13.5, peaking at E15.5, and declining thereafter until P0 (Fig1A). MyHC-slow exhibited a similar transcript expression profile as MyHC-emb, starting to rise at E10.5, peaking at E15.5, and declining thereafter (Fig1A′). A striking difference in magnitude of the transcript relative expression profiles was observed, with MyHC-emb reaching a relative fold change of about 6000, while MyHC-slow recorded a peak relative fold change of about 150 (Fig1A,A′). At the protein level, MyHC-emb and -slow were expressed at E10.5 in the somitic region with relatively lower levels of MyHC-slow as compared to MyHC-emb (Fig1B, B′, C, C′). By E13.5, during embryonic myogenesis, both MyHC-emb and -slow were expressed in hind limb muscles, with all muscle fibers expressing both MyHCs (Fig1D, D′, E, E′). By E16.5, during fetal myogenesis, MyHC-slow is expressed in a subset of myofibers compared to MyHC-emb in the hind limb (Fig1F, F′, G, G′). MyHC-emb protein levels in the hind limb relative to E12.5 expression peaks at two time points, a minor one at E14.5 and a major one at E16.5, compared to MyHC-slow which is expressed at relatively constant levels until E16.5, peaking at E17.5 (Fig1H). During C2C12 myogenic cell differentiation in vitro, both MyHC-emb and -slow transcripts were detectable by qPCR from day2 of differentiation onwards, with transcript levels peaking at day6-7 of differentiation and falling thereafter (Fig1I, J). As was observed in vivo, there was considerable difference in the magnitude of transcript expression profiles between MyHC-emb and -slow, with MyHC-emb reaching relative fold changes of close to 36,000, while MyHC-slow recorded a peak relative fold change of about 12,000 (Fig1I, J). MyHC-emb protein was detected from day 4 of C2C12 differentiation by western blots, peaking at day 8 and decreasing thereafter (Fig1K, L). MyHC-slow protein was detected from day 5 of C2C12 differentiation, increasing gradually until day 13 (Fig1K, M).

**Figure 1:**
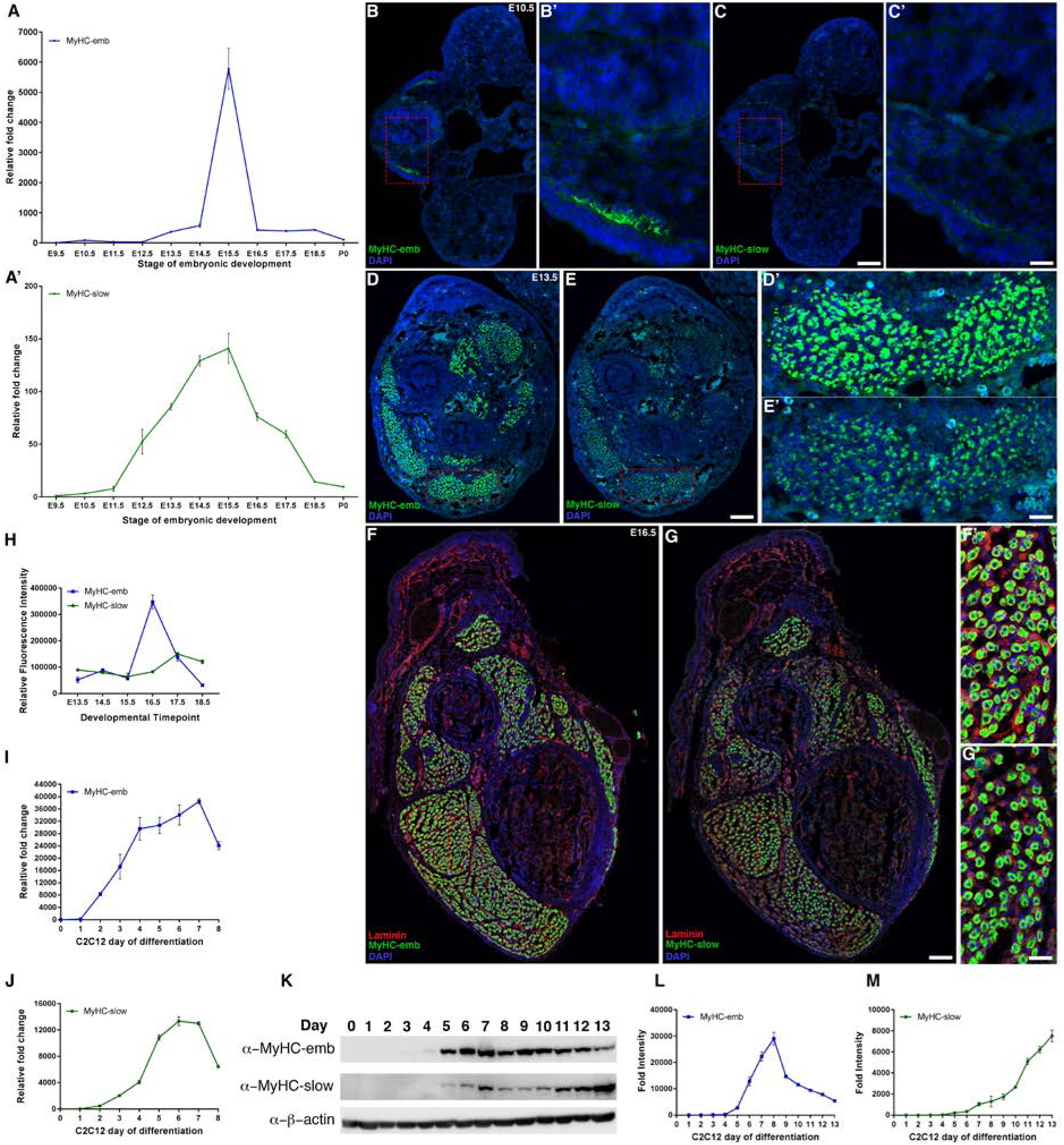
MyHC-embryonic and -slow are expressed early during mouse embryonic development and C2C12 myogenic differentiation. MyHC-emb (A) and MyHC-slow (A') RNA expression detected by qPCR, starts at around E10.5, increasing gradually and peaking at E15.5, declining thereafter until birth (note the Y-axis scale difference between MyHC-emb and -slow, indicating that MyHC-emb transcripts are expressed many folds higher than MyHC-slow). The transcript levels for both MyHC-emb and -slow were normalized to E9.5 expression. MyHC-emb (B, B') and -slow (C, C') proteins are expressed in the somitic region (boxed region in B and C are magnified in B' and C' respectively) in the mouse embryo at E10.5, in the limb muscle fibers during embryonic myogenesis at E13.5 (D, D', E, E', where D' and E' magnifications of the boxed region in D and E respectively), and in the limb muscle fibers during fetal myogenesis at E16.5 (F, F', G, G', with F' an G' magnifications of the boxed region in F and G respectively). The protein expression levels of MyHC-emb and MyHC-slow were calculated over the course of development by analyzing the fluorescence intensity relative to E12.5, with MyHC-emb exhibiting a minor peak at E14.5 and a major peak at E16.5, declining thereafter, whereas MyHC-slow levels were relatively constant until E16.5, peaking at E17.5 following which it started declining (H). During C2C12 mouse myogenic cell differentiation, MyHC-emb (I) and MyHC-slow (J) RNA expression detected by qPCR increases as differentiation progresses, with MyHC-emb levels peaking by day 7 of differentiation and MyHC-slow levels peaking by day 6 of differentiation (note the Y-axis scale difference between MyHC-emb and -slow, indicating that MyHC-emb transcripts are expressed many folds higher than MyHC-slow). MyHC-emb (K, L) and -slow (K, M) proteins are expressed by day 5 of C2C12 cell differentiation, with MyHC-emb levels peaking by day 7 (L), and MyHC-slow levels increasing until the thirteenth day of differentiation (M). (Scale bars in C, E, and G is 100 microns; scale bars in C’, E’, and G' is 25 microns).

Overall, we found that MyHC-emb is expressed at high levels in all embryonic myofibers and suggests that MyHC-emb might play an essential role in embryonic myogenesis (Beylkin, et al., 2006; Lu, et al., 1999).

### *MyH3^fl3-7^* and *MyH3^Δ^* alleles allow in vivo genetic deletion of MyHC-embryonic

In order to test whether MyHC-emb is required for myogenic differentiation in vivo, we generated a conditional *MyH3^fl3-7^* allele for MyHC-emb in mice, where exons 3-7 of MyHC-emb were flanked by LoxP sites (Fig2A). By crossing the *MyH3^fl3-7^* allele to the germline deleter *Hprt^Cre^* driver, we generated the *MyH3^Δ^* allele, where exons 3-7 were deleted (Fig2B). Both *MyH3^fl3-7^* and *MyH3^Δ^* alleles were verified by PCR (Supplementary Fig1A, B, C). In crosses between *MyH3^Δ/+^* heterozygous mice, *MyH3^Δ/Δ^* homozygous pups were not obtained at the expected 3:1 ratio, but at about 7:1 ratio (17 out of 126), suggesting that approximately half of the animals of the *MyH3^Δ/Δ^* genotype die in utero during embryonic or fetal stages, and that MyHC-emb might have critical functions during developmental stages. We also validated that *MyH3^Δ/Δ^* animals are null for MyHC-emb by qPCR for MyHC-emb using cDNA derived from multiple muscles of *MyH3^Δ/Δ^* mice (Fig2C). At the protein level, MyHC-emb could not be detected by immunofluorescence using an antibody specific to MyHC-emb on cross sections through the shank region of *MyH3^Δ/Δ^* mice (Fig2D, D′ to E, E′).

### Loss of MyHC-embryonic results in neonatal mis-regulation of other MyHC isoforms in vivo

Since MyHC-emb is tightly linked with other MyHC isoforms, we tested whether loss of MyHC-emb expression leads to mis-regulation of other MyHC isoforms (Mathew, et al., 2011; Allen and Leinwand, 2001; Sartorius, et al., 1998; Gunning and Hardeman, 1991). MyHC distribution within skeletal muscles differs according to their location and function and therefore we tested 4 different muscles, tibialis anterior, quadriceps, gastrocnemius, and diaphragm, with varying amounts of the different MyHC isoforms. We found that in all 4 muscles, MyHC-IIa isoform transcript levels were significantly upregulated in *MyH3^Δ/Δ^* animals (Fig2G, H, I, J). MyHC-IIx transcript levels were significantly upregulated in the quadriceps and diaphragm of *MyH3^Δ/Δ^* mice (Fig2H, J), while MyHC-peri was also significantly upregulated in the quadriceps and gastrocnemius (Fig2H, I). Remarkably, MyHC-IIa, which is upregulated in all of the muscles, is located adjacent to MyHC-emb in the MyHC cluster on chromosome 11, while MyHC-IIx and MyHC-peri are located downstream of MyHC-IIa (Weiss, et al., 1999; Vikstrom, et al., 1997) (Fig 2F). Thus, loss of MyHC-emb leads to compensatory upregulation of other MyHCs in the fast MyHC gene cluster, with the genes located closest to MyHC-emb physically upregulated in more muscles. MyHC-slow, which is not part of the MyHC chromosome 11 cluster, transcript levels were significantly downregulated in *MyH3^Δ/Δ^* animals in the tibialis anterior and quadriceps (Fig2G, H). An intriguing aspect of these findings is that although MyHC-emb is expressed in all muscles during development, the effect of loss of MyHC-emb is not uniform, with different effects on different muscles.

**Figure 2:**
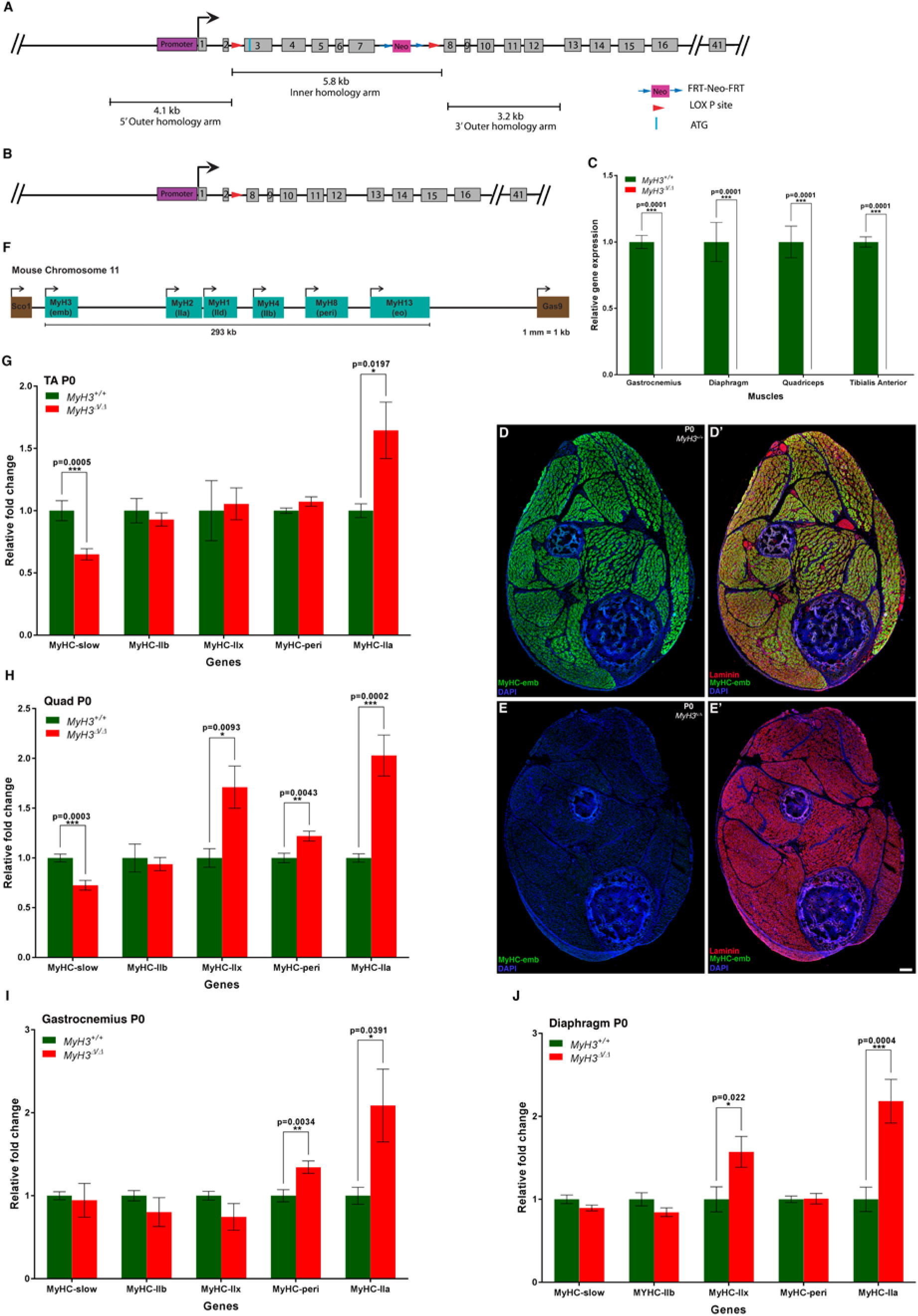
MyH3 targeted alleles allow for genetic manipulation of MyHC-embryonic and loss of MyHC-embryonic leads to neonatal mis-regulation of other MyHC isoforms. Schematics depicting *MyH3^fl3-7^* conditional allele, where LoxP sites (red arrowheads) flank exons 3-7 to facilitate conditional deletion of this genomic region (A) and the *MyH3^Δ^* allele that was generated by crossing *MyH3^fl3-7^* animals to a germline deleter *Hprt^Cre^* driver, following which the animals lack exons 3-7 of MyHC-emb (B). *MyH3^Δ/Δ^* is null for MyHC-emb at the transcript level by qPCR, where there was no detectable MyHC-emb transcripts in the gastrocnemius, diaphragm, quadriceps and tibialis anterior muscles, compared to control *MyH3^+/+^* animals at P0 (C). MyHC-emb protein was undetectable by immunofluorescence in cross sections through *MyH3^Δ/Δ^* P0 hind limbs (E, E′), as compared to control *MyH3^+/+^* animals (D, D′). Schematic showing the MyHC gene cluster on mouse chromosome 11 which is approximately 293 kb in size, where MyHC-emb is followed by MyHC-IIa, -IId, -IIb, -perinatal and -extraocular (F). In P0 neonates, loss of MyHC-emb in *MyH3^Δ/Δ^* animals leads to a significant upregulation in the transcript levels of MyHC-IIa in all 4 muscles studied, namely tibialis anterior, quadriceps, gastrocnemius and diaphragm (G, H, I and J respectively) compared to *MyH3^+/+^* animals by qPCR; transcript levels of MyHC-IIx are significantly upregulated in the quadriceps and diaphragm muscles in *MyH3^Δ/Δ^* animals (H and J), and MyHC-perinatal are significantly upregulated in the quadriceps and gastrocnemius muscles in *MyH3^Δ/Δ^* animals by qPCR (H and I); transcript levels of MyHC-slow are significantly down-regulated in the tibialis anterior and quadriceps muscles in *MyH3^Δ/Δ^* animals by qPCR (G and H). (Scale bar in E' is 100 microns).

### Loss of MyHC-embryonic leads to alterations in myofiber number, area and fiber type in vivo

Next, we investigated whether loss of MyHC-emb function causes myofiber differentiation defects in vivo since previous studies on MyHC-IIb and IIx null adult mice report effects on myofiber size, and structure (Sartorius, et al., 1998; Acakpo-Satchivi, et al., 1997). For this, we quantified the number of myofibers in the EDL (extensor digitorum longus) and soleus muscles, which are muscles rich in fast and slow fibers respectively, at P0. We found that there was a significant increase in the total number of myofibers in the EDL upon loss of MyHC-emb, while there was no difference in the soleus (Fig3A, B). This indicates that MyHC-emb is required to regulate myofiber number in muscles rich in fast fibers whereas it does not play this role in slow muscles. We next tested whether loss of MyHC-emb affected muscle fiber type. In order to analyze the number of MyHC-slow+ fibers, we labeled *MyH3^+/+^* and *MyH3^Δ/Δ^* P0 mouse hind limb cross sections through the shank with MyHC-slow and laminin to mark the basal lamina (Fig3C, C’, C’’ and D, D’, D’’ respectively, with C’’ and D’’ magnified regions indicated by the red rectangles in C’ and D’). We found a striking increase in the number of MyHC-slow+ fibers in the *MyH3^Δ/Δ^* animals (Fig3C, C’, C’’, D, D’, D’’, and G). We also observed that the size of the myofibers was reduced in the soleus muscle upon loss of MyHC-emb (Fig3E, F). We quantified the myofiber area of the soleus and EDL muscles of *MyH3^+/+^* and *MyH3^Δ/Δ^* P0 mice by grouping myofibers in 0-50 μm^2^, 50-150 μm^2^, and above 150 μm^2^ categories (Fig3H, I). We found that there was a significant increase in the myofibers with the smallest area (0-50 μm^2^) in the soleus upon loss of MyHC-emb, while there was no significant difference in any of the myofiber area categories upon loss of MyHC-emb in the EDL (Fig 3H, I). Thus, MyHC-emb regulates myofiber cross-sectional area in muscles rich in slow fibers but not in fast fiber rich muscles.

**Figure 3:**
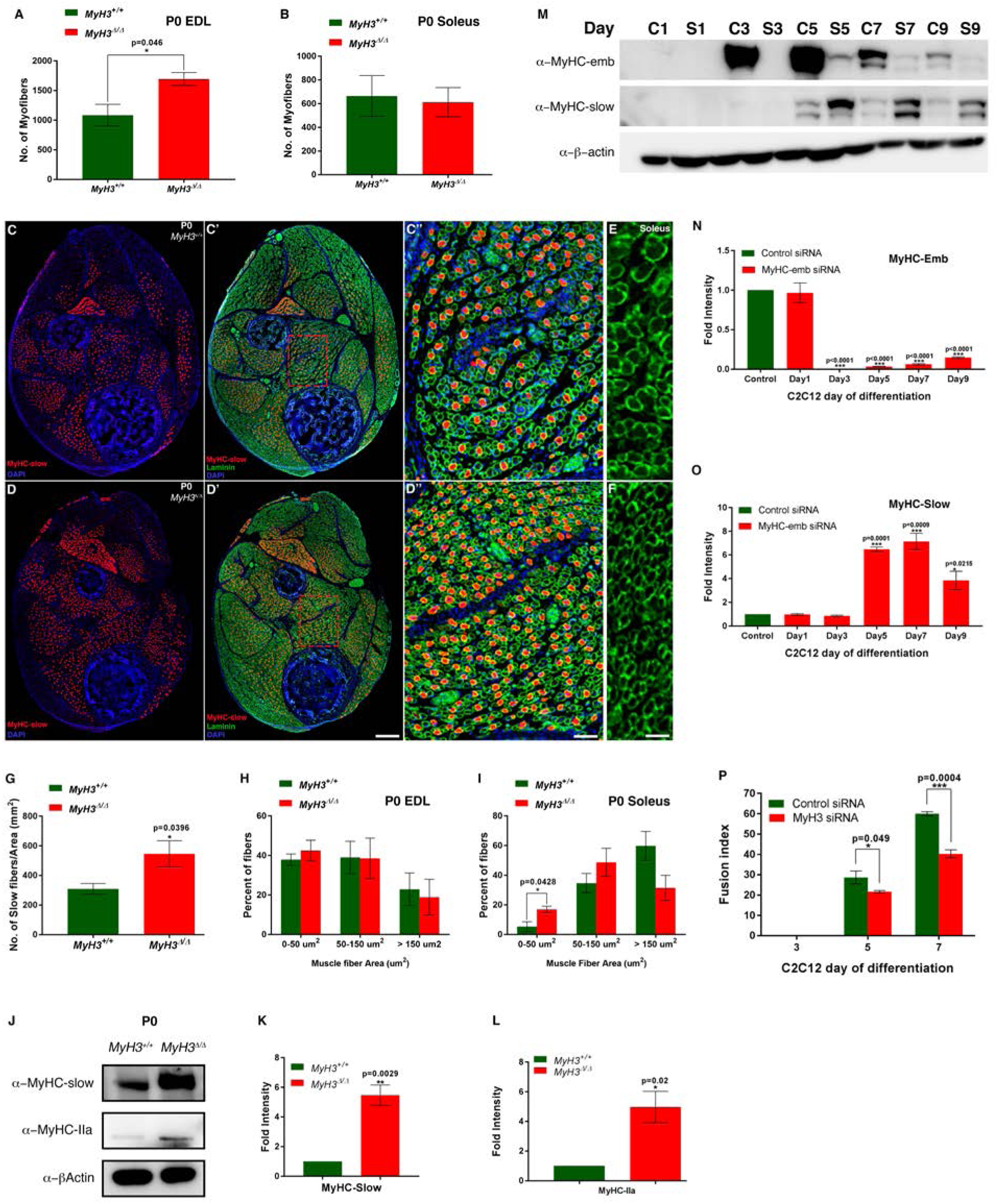
Loss of MyHC-embryonic function leads to myogenic differentiation defects in vivo and vitro. At postnatal day 0, the number of myofibers are significantly increased in the EDL muscle in the hind limbs of *MyH3^Δ/Δ^* animals compared to *MyH3*^+/+^ animals (A), while no significant difference in the number of myofibers is observed in the case of the soleus muscle (B). MyHC-slow positive fibers are increased in P0 *MyH3^Δ/Δ^* animal hind limb cross sections (D, D’), compared to *MyH3^+/+^* control animals (C, C’), where the sections are labeled by immunofluorescence for MyHC-slow (Red), Laminin (Green) and DAPI (Blue). C’’ and D’’ are magnified regions of the boxed area of *MyH3^+/+^* and *MyH3^Δ/Δ^* P0 hind limb cross sections from C’ and D’ respectively. The elevated number of MyHC-slow+ fibers in P0 *MyH3^Δ/Δ^* hind limb cross sections compared to *MyH3^+/+^* controls was quantified normalized to total area and is statistically significant (G). The myofiber size was found to be smaller in the soleus muscle of *MyH3^Δ/Δ^* animals (F) as compared to *MyH3^+/+^* animals (E). The myofiber area of myofibers of the EDL and soleus muscles at P0 were quantified, grouping myofibers into 3 categories by area, 0-50 μm^2^, 50-150 μm^2^and above 150 μm^2^. It was found that the number of myofibers in the 0-50 μm^2^group was increased significantly in the soleus muscle of *MyH3^Δ/Δ^* animals compared to controls (I), but was unchanged in the EDL (H). Protein levels of MyHC-slow and MyHC-IIa are about 5-fold upregulated in the *MyH3^Δ/Δ^* animals compared to *MyH3^+/+^* animals by western blots (J), which is quantified by densitometry (K and L). MyHC-emb protein levels are significantly reduced upon treatment of C2C12 cells with *MyH3*siRNA compared to control siRNA over 9 days of differentiation as detected by western blots and densitometry (M and N), while protein levels of MyHC-slow are significantly increased from day 5 onwards reaching a peak of more than 6-fold increase by day 7 upon MyHC-emb knockdown as detected by western blots and densitometry (M and O). Knockdown of MyHC-emb results in a significantly reduced fusion index at days 5 and 7 of C2C12 differentiation, compared to control siRNA treated cells (P). (Scale bar in D′ is 200 microns, D′′ is 40 microns and F is 20 microns respectively).

Since we observed an increase in the number of MyHC-slow+ fibers upon loss of MyHC-emb, we validated the increase in MyHC-slow protein levels by western blot, and found a significant 6-fold increase in MyHC-slow protein levels in MyHC-emb null P0 muscle lysates (Fig 3J, K). In addition, we also observed a significant 5-fold increase in MyHC-IIa protein levels upon loss of MyHC-emb (Fig3J, L). Interestingly, at the transcript level, we had observed a down-regulation of MyHC-slow upon loss of MyHC-emb in the tibialis anterior and quadriceps muscles (Fig2G, H) whereas the number of MyHC-slow+ fibers and MyHC-slow protein levels were significantly increased upon loss of MyHC-emb, indicating that post-transcriptional or protein stability mechanisms are involved in regulating the levels of MyHC-slow (Fig3G, K).

We also investigated the effect of MyHC-emb knockdown by siRNA during C2C12 myogenic differentiation in vitro. The knockdown was verified at the protein level by western blots for MyHC-emb, and 80% or higher knockdown efficiency was observed (Fig3M, N). As was the case in vivo, protein levels of MyHC-slow exhibited a significant ∼7-fold increase upon MyHC-emb knockdown at days 5 and 7 of differentiation (Fig3M, O). We also found that differentiation was compromised upon MyHC-emb knockdown, with the fusion index significantly lower in *MyH3*siRNA treated cells on days 5 and 7 of differentiation (Fig3P). In summary, these results indicate that MyHC-emb is important for proper myogenic differentiation, regulating fiber number, fiber size, fiber type and fusion index.

### Distinct muscles respond differently to loss of MyHC-embryonic

Since we observed that loss of MyHC-emb led to alterations in myofiber characteristics, we tested whether loss of MyHC-emb had other effects on myogenic differentiation. Therefore, we carried out a whole-transcriptome mRNA-sequencing (RNA-Seq) experiment comparing gene expression between P0 *MyH3^+/+^* and *MyH3^Δ/Δ^* animals, using tibialis anterior, quadriceps, gastrocnemius, and diaphragm muscles, which express varying amounts of the different MyHC isoforms (Fig4A, B, C, D). We found 218 genes were significantly mis-regulated in MyHC-emb null muscles (Supplementary Table 3). Only 2 genes, Nfil3 and Btg2, were significantly mis-regulated across all 4 muscles, while 18 genes were significantly mis-regulated across any 3 muscles, and 42 genes significantly mis-regulated across 2 muscles (Fig4H and Supplementary Table 3).

**Figure 4:**
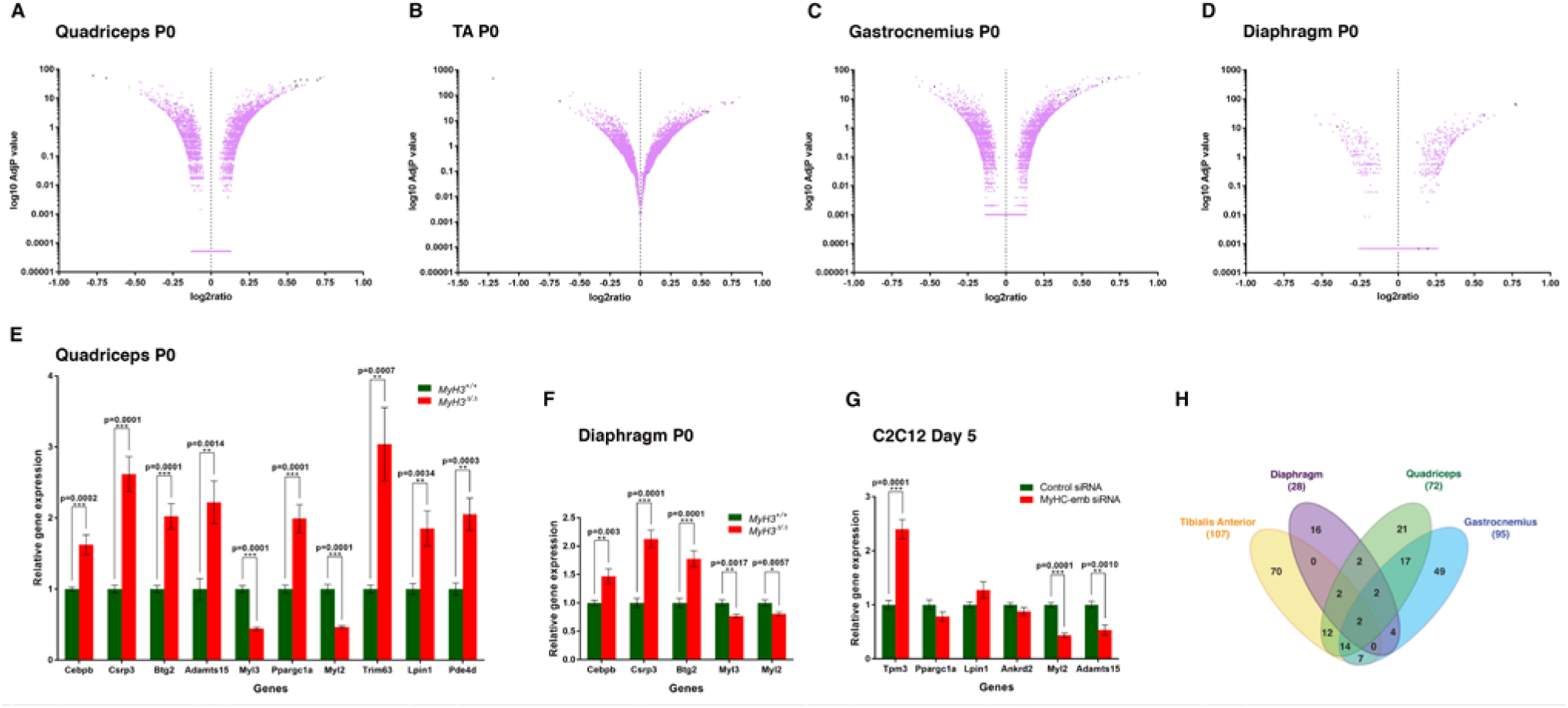
Loss of MyHC-embryonic leads to global mis-regulation of genes involved in myogenic differentiation. Volcano plots depicting results from RNA sequencing experiment comparing P0 *MyH3^+/+^* muscle samples to *MyH3^Δ/Δ^* samples for quadriceps, tibialis anterior, gastrocnemius, and diaphragm muscles respectively (A-D). The adjusted P values are on a log10 scale, and candidates that are significantly up or down regulated are marked as dark spots on the volcano plot (A-D). Selected candidate genes from the RNA sequencing experiment were validated for up or down regulation by qPCR on *MyH3^+/+^* samples compared to *MyH3^Δ/Δ^* samples for the P0 quadriceps muscle (E) or P0 diaphragm muscle (F). 6 candidates from the RNA sequencing experiment were tested for mis-regulation in day 5 *MyH3* siRNA treated C2C12 cells compared to control siRNA treated cells, and 3 were found to be significantly mis-regulated (G). Venn diagram depicting number of candidates obtained and degree of overlap in the RNA sequencing results comparing P0 *MyH3^+/+^* muscle samples to *MyH3^Δ/Δ^* samples for quadriceps, tibialis anterior, gastrocnemius, and diaphragm muscles (H).

We found that the genes mis-regulated in 2 or more muscles in the RNA-Seq experiment were involved in myogenic differentiation. For instance, Btg2, a transcriptional co-regulator, which is mis-regulated in all 4 muscles, is involved in muscle development and differentiation (Feng, et al., 2007) and Adamts15, a metalloproteinase mis-regulated in 3 muscles, is essential for myoblast fusion (Stupka, et al., 2013). Csrp3, a muscle LIM domain protein mis-regulated in 3 muscles, interacts with several myogenic regulatory proteins and functions in myofibril organization and contractile apparatus maintenance (Arber, et al., 1997; Kong, et al., 1997), interacting with sarcomeric Z-disc proteins such as Zyxin, α-actinin (Louis, et al., 1997), acting as a mechanical stress sensor (Wang, et al., 2006) to roles in muscle autophagy (Rashid, et al., 2015). Transcripts for genes which encode proteins essential for muscle contractility such as Tpm3, Myl3 and Myl2 were mis-regulated in multiple muscles (Supplementary Table 3). 12 genes mis-regulated upon loss of MyHC-emb were also identified in a study to discover transcriptional regulators involved in early muscle differentiation using C2C12 myoblasts (Rajan, et al., 2012) (Supplementary Table 4). Of these 12 candidates, 8 genes were mis-regulated in more than one muscle in our study (Supplementary Table 4). 6 of these candidates were tested for their role in early C2C12 differentiation by knockdown assays and found to be vital for differentiation by Rajan et al (Supplementary Table 4). All of this validates that MyHC-emb is necessary for normal myogenic differentiation, and loss of MyHC-emb leads to mis-regulation of muscle differentiation-related genes.

Overall, the diaphragm shared the least number of mis-regulated candidates as compared to the 3 limb muscles, possibly because the diaphragm is a specialized muscle that is anatomically and functionally distinct from the limb (Fig4H). 156 genes were uniquely mis-regulated in a single muscle type, which could be genes that have distinctive roles in those specific muscles, depending on the fiber type characteristics, functional properties and anatomical position (Fig4H). Some of the MyHCs that were tested for mis-regulation by qPCR were also found to be mis-regulated in this experiment, for instance MyHC-IIa was upregulated in the gastrocnemius, and MyHC-slow was down-regulated in the tibialis anterior (Supplementary Table 3).

Transcripts of Trim63 (MuRF1), a muscle specific ubiquitin ligase that targets proteins for degradation (Witt, et al., 2005), is upregulated in *MyH3* null animals, which could explain why they exhibit increased cell death, marked by elevated Caspase-3 levels (Fig5A, G) and (Supplementary Table 3).

**Figure 5:**
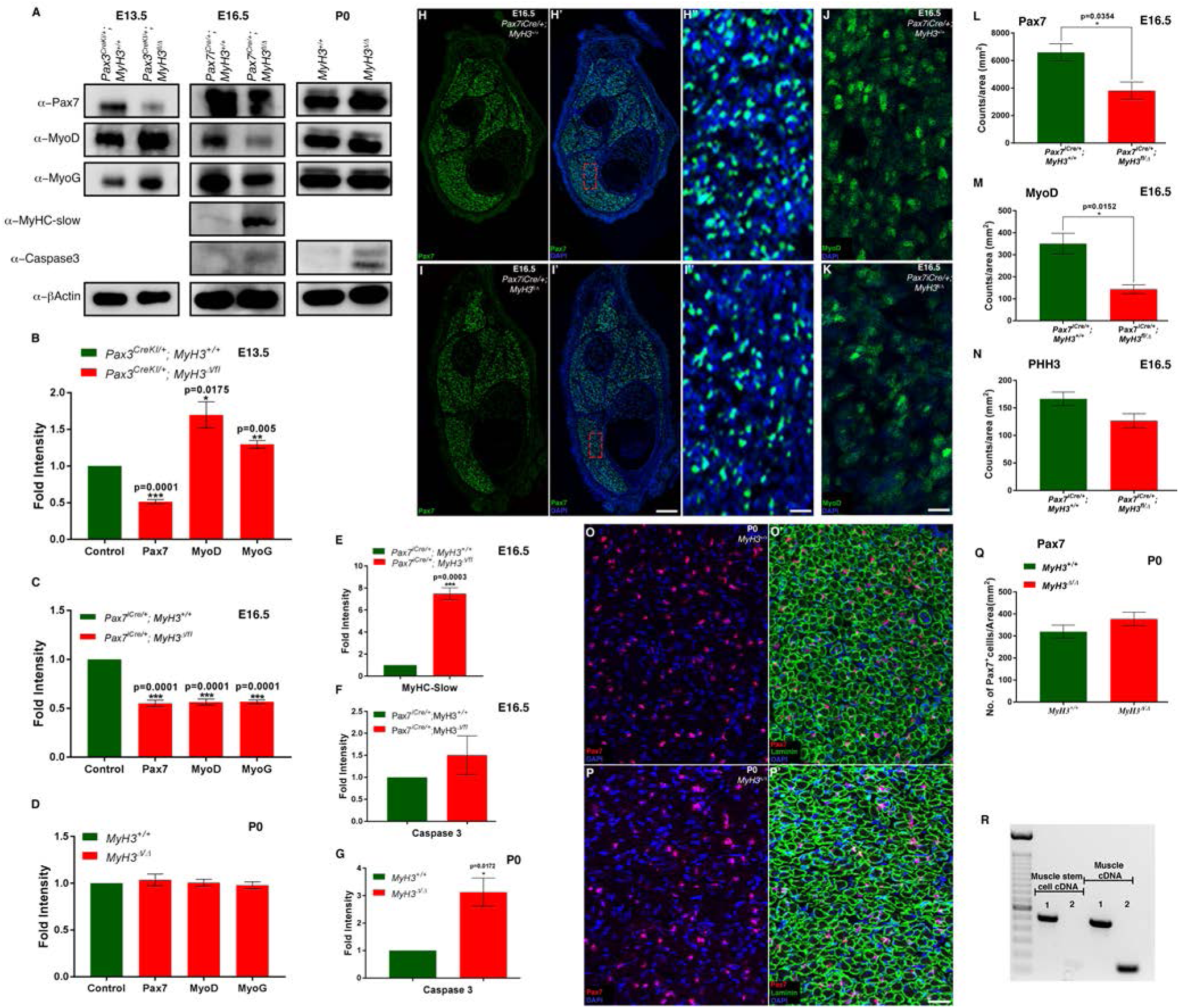
MyHC-embryonic regulates myogenic progenitor and myoblast differentiation non-cell autonomously during embryonic and fetal phases of myogenesis in vivo. At E13.5 during embryonic myogenesis, loss of MyHC-emb in the myogenic lineage in *Pax3^CreKI/+^, MyH3Δ/fl3-7* leads to a significant reduction in Pax7 levels and a significant increase in MyoD and Myogenin levels by western blots (A), and quantified by densitometry (B) compared to control *Pax3^CreKI/+^, MyH3+/+* animals. At E16.5 during fetal myogenesis, loss of MyHC-emb in the myogenic lineage in *Pax7^iCre/+^, MyH3Δ/fl3-7* leads to a significant reduction in Pax7, MyoD and Myogenin levels by western blots (A), and quantified by densitometry (C), as compared to control *Pax7^iCre/+^, MyH3+/+* animals. Pax7, MyoD and Myogenin protein levels are similar between *MyH3^Δ/Δ^* and *MyH3^+/+^* animals at P0 by western blots (A), which is quantified by densitometry (D). The levels of MyHC-slow are increased about 7-fold at E16.5 during fetal myogenesis compared to controls in *Pax7^iCre/+^, MyH3Δ/fl3-7* animals by western blot (A) and quantified by densitometry (E). Cell death measured by levels of cleaved Caspase3 protein was detected by western blots at E16.5 and P0 (A) and quantified by densitometry which showed that there was no significant difference in cell death between controls and *Pax7^iCre/+^, MyH3Δ/fl3-7* animals at E16.5 (F), while cell death was significantly elevated by about 3-fold in the *MyH3^Δ/Δ^* animals compared to *MyH3^+/+^* animals at P0 (G). At E16.5 during fetal myogenesis, loss of MyHC-emb in the myogenic lineage in *Pax7^iCre/+^, MyH3Δ/fl3-7* leads to a significant reduction in the number of Pax7+ cells in the hind limb muscles compared to control by labeling Pax7 (Green), and DAPI (Blue) by immunofluorescence (H, H’, I, I’), which is quantified normalized to total area (L). H′’ and I′’ are magnified regions from H’ and I’ respectively. MyoD+ myoblasts at E16.5 during fetal myogenesis in control and *Pax7^iCre/+^, MyH3Δ/fl3-7* embryo hind limb muscles where MyoD (Green) and DAPI (Blue) were labeled by immunofluorescence (J, K) and a significant reduction in the number of MyoD+ myoblasts was observed in the *Pax7^iCre/+^, MyH3Δ/fl3-7* embryos, which is quantified normalized to total area (M). No significant difference in the number of dividing, phospho-histone H3+ cells were observed at E16.5 in *Pax7^iCre/+^, MyH3Δ/fl3-7* embryo hind limb muscles compared to controls, which is quantified normalized to total area (N). There was no significant difference in the number of Pax7+ cells by immunofluorescence between *MyH3^+/+^* control and *MyH3^Δ/Δ^* knockout P0 hind limb cross sections (O-P’), where Pax7 (Red), Laminin (Green) and DAPI (Blue) have been labeled. The number of Pax7+ cells in P0 *MyH3^Δ/Δ^* hind limb cross sections compared to *MyH3^+/+^* controls was quantified normalized to total area and no significant difference in numbers were observed (Q). MyHC-emb transcripts are not detectable in muscle stem cell cDNA as opposed to muscle cDNA by semi-quantitative RT-PCR (R); 1 represents PCR for a positive control (GAPDH) and 2 represents PCR for MyHC-emb (R). (Scale bar in I’ is 200 microns, and in I'' and K are 20 microns).

To validate the the RNA-Seq results, we carried out qPCR to detect levels of selected candidate genes in the quadriceps and diaphragm P0 *MyH3^+/+^* and *MyH3^Δ/Δ^* muscles (Fig4E, F). We found that 10 genes in the case of the quadriceps and 5 genes in the case of the diaphragm, were mis-regulated as was observed in the RNA-Seq, validating the RNA-Seq results (Fig4E, F). We also found that 3 out of 6 candidates from the RNA-Seq experiment that we tested were mis-regulated upon MyHC-emb knockdown during the course of C2C12 myogenic differentiation, indicating that MyHC-emb affects similar processes and pathways in vivo and in vitro (Fig4G).

In summary, genes related to differentiation were mis-regulated upon loss of MyHC-emb function, indicating that MyHC-emb is critical for proper myogenic differentiation. Although MyHC-emb is expressed uniformly across muscles, the mis-regulated genes are not the same in different muscles, possibly due to inherent differences between muscles, reflecting their fiber type and metabolic diversity.

### MyHC-embryonic non-cell autonomously regulates myogenic progenitor differentiation during embryonic and fetal myogenesis

Since we observed myogenic differentiation defects in muscles of *MyH3^Δ/Δ^* animals at neonatal stages, and since MyHC-emb starts to be downregulated by P0, we hypothesized that the differentiation defects are most likely due to embryonic or fetal specific requirement of MyHC-emb. To test the embryonic and fetal specific roles played by MyHC-emb, we made use of 2 Cre drivers: *Pax3^CreKI/+^* which causes Cre mediated recombination in embryonic and fetal myogenic lineages and *Pax7^iCre/+^* which recombines only in the fetal myogenic lineages (Hutcheson, et al., 2009; Engleka, et al., 2005; Keller, et al., 2004). At E13.5, during embryonic myogenesis, deletion of MyHC-emb in *Pax3^CreKI/+^, MyH3Δ/fl3-7* led to a significant reduction of the myogenic progenitor marker Pax7 protein levels, to half as that of control *Pax3^CreKI/+^, MyH3+/+* embryos (Fig5A, B), while protein levels of the committed myoblast markers MyoD and Myogenin were significantly increased (Fig5A, B). This suggests that loss of MyHC-emb accelerates the differentiation of myogenic progenitors leading to the depletion of the progenitors as seen by a reduction in Pax7 levels, and a concomitant increase in the differentiated myoblasts as indicated by the increased levels of the committed myoblast markers MyoD and Myogenin (Fig5A, B).

At E16.5, during fetal myogenesis, deletion of MyHC-emb in *Pax7^iCre/+^, MyH3Δ/fl3-7* also resulted in a significant reduction in the levels of Pax7 protein (Fig5A, C). Interestingly, the levels of the committed myoblast markers MyoD and Myogenin were also significantly decreased in the E16.5 *Pax7^iCre/+^, MyH3Δ/fl3-7* embryos (Fig5A, C). MyHC-slow protein levels were upregulated by about 7-fold in the E16.5 *Pax7^iCre/+^, MyH3Δ/fl3-7* embryos compared to control *Pax7^iCre/+^, MyH3+/+* embryos (Fig5A, E). This suggests that the high levels of MyHC-slow protein and increased number of MyHC-slow+ fibers seen at P0 (Fig3G, J, K) could be due to increased MyHC-slow protein levels during fetal myogenesis in the absence of MyHC-emb.

We investigated this further at neonatal stages by western blots using protein lysates from *MyH3^+/+^* and *MyH3^Δ/Δ^* P0 mouse hind limb muscles and quantified the levels of Pax7 by densitometry which indicated that there was no difference in Pax7 levels between control and MyHC-emb null animals (Fig5A, D). In order to test whether there were alterations in the number of differentiating myoblasts or protein levels of MRFs that facilitate myogenic differentiation, we also performed western blots for MyoD and Myogenin, and no significant difference in their protein levels were observed upon densitometry (Fig5A, D).

The observed differences in the levels of myogenic progenitor and myoblast markers could be due to alterations in cell death. We tested this by performing western blots for cleaved caspase 3 at E16.5 where MyHC-emb was deleted during fetal myogenesis and at P0. No significant differences in cleaved caspase 3 levels between MyHC-emb deleted and control protein lysates were observed at E16.5, but an approximately 3-fold increase in cell death was seen in the MyHC-emb null P0 muscle lysates compared to controls (Fig5A, F, G). Thus, alterations in cell death is not responsible for the observed differences in myogenic progenitor and myoblast marker protein levels during fetal myogenesis at E16.5, upon loss of MyHC-emb function. The increased cell death at P0 could be due to aberrant differentiation in the *MyH3^Δ/Δ^* animals, and fits well with increased Trim63 (MuRF1) levels, a muscle specific ubiquitin ligase that targets proteins for degradation, in P0 *MyH3^Δ/Δ^* muscles by RNA-Seq (Witt, et al., 2005).

As with embryonic myogenesis, loss of MyHC-emb could be accelerating the differentiation of myogenic progenitors during fetal myogenesis, leading to the depletion of the progenitors as seen by the reduction in Pax7 levels. To verify this, we examined whether there was any difference in the number of Pax7+ muscle progenitors between *Pax7^iCre/+^, MyH3+/+* control and *Pax7^iCre/+^, MyH3Δ/fl3-7* MyHC-emb deleted embryos, by labeling Pax7 and DAPI by immunofluorescence in E16.5 mouse hind limb cross sections through the shank (Fig5H, H’, H′′ and I, I’, I′′ with H′′ and I′′ the magnified regions marked by the rectangle in H′ and I′ respectively). We found that there was a 50% reduction in the number of Pax7+ myogenic progenitors in the *Pax7^iCre/+^, MyH3Δ/fl3-7* embryos (Fig5H′′, I′′, L). Similarly, there was a statistically significant decline of more than 60% MyoD+ myoblasts in the *Pax7^iCre/+^, MyH3Δ/fl3-7* embryos (Fig5M). We did not observe any significant difference in the number of dividing, phospho-histone H3+ nuclei between *Pax7^iCre/+^, MyH3Δ/fl3-7* embryos, indicating that rate of cell division is not responsible for the observed decrease in Pax7+ progenitors and MyoD+ myoblasts (Fig5N). This suggests that loss of MyHC-emb accelerates the differentiation of myogenic progenitors during fetal myogenesis and leads to a depletion of Pax7+ progenitors and MyoD+ myoblasts. We examined whether there were any differences in the number of Pax7+ muscle progenitors between *MyH3^+/+^* and *MyH3^Δ/Δ^* P0 mice and found no significant difference (Fig5O, O’, P, P’ and Q).

Since MyHC-emb seems to be involved in myogenic differentiation in vivo, it is crucial to demonstrate which cell types express MyHC-emb. Therefore, although it has been well accepted that MyHC-emb transcript is only expressed by differentiated muscle fibers and not by muscle progenitor populations, we performed semi-quantitative PCR using MyHC-emb specific primers on P0 cDNA derived from muscle progenitors compared to cDNA derived from whole muscle where the bulk of the RNA will be myofiber derived. We observed that the MyHC-emb specific PCR amplicon was obtained only with the whole muscle cDNA and not with the muscle progenitor cDNA, thus validating that MyHC-emb is not expressed in undifferentiated myogenic cell types, and is only expressed by terminally differentiated myofibers (Fig5R). This indicates that the effect of MyHC-emb on differentiation, especially on progenitor cells and myoblasts has to be mediated by non-cell autonomous signals arising from the myofibers which express MyHC-emb.

In summary, MyHC-emb is essential for embryonic and fetal myogenesis by non-cell autonomously regulating the differentiation rate of myogenic progenitors and myoblasts. This role of MyHC-emb is independent of cell death or cell proliferation and is restricted to developmental stages. By neonatal stages, MyHC-emb is not required presumably because other MyHC isoforms such as MyHC-IIa are able to compensate the loss of MyHC-emb. Loss of MyHC-emb also resulted in an increase in MyHC-slow protein levels during fetal myogenesis, which might explain the drastic increase in the number of MyHC-slow+ fibers observed at P0.

### MyHC-embryonic is required for proper myogenesis in vitro

Next, we wished to test whether loss of MyHC-emb function had similar effects in vitro, using C2C12 myogenic cells. In order to investigate whether loss of MyHC-emb leads to an altered rate of C2C12 differentiation, we looked at the reserve cell pool, which is the population of undifferentiated cells, by labeling for F-actin using phalloidin, where the mono-nuclear reserve cells appeared brightly stained (Ricotti, et al., 2011; Burattini, et al., 2004) (Fig6A, A’, B, B’). We observed that the number of reserve cells were reduced in the *MyH3* siRNA treated C2C12 cells at day 5 of differentiation as compared to control siRNA treated cells (white arrows in B’ and A’ respectively). We repeated these experiments to quantify the number of reserve cells normalized to unit area, and found that there was a significant, greater than 6-fold reduction in the number of reserve cells in the *MyH3* siRNA treated cells (Fig6C). We hypothesized that any of three possibilities could explain the reduction in reserve cells: reduced cell proliferation, increased cell death, or increased rate of differentiation leading to their depletion. We found no change in number or proliferative phospho-histone H3+ cells between control and *MyH3* siRNA treated cells (Fig6D). In order to quantify levels of cell death, we performed western blots for cleaved caspase3, and found that although there was a significant increase in cell death on day 1 of MyHC-emb knockdown, the cell death was significantly less compared to control samples on days 5, 7 and 9 of differentiation (Fig6E, H). These results indicate that it is unlikely that the reduction in the number of reserve cells observed upon MyHC-emb knockdown is due to decreased rate of cell division or increased cell death, and must be due to alteration in the rate of myogenic differentiation. We tested this by analyzing the protein expression of the MRFs MyoD and Myogenin following MyHC-emb knockdown, where we observed an initial upregulation of expression compared to control cells on days 3-5 of differentiation (except for Myogenin at day 3, where we observed a downregulation), followed by a significant reduction in expression of both at later time points, days 7-9 of differentiation (Fig6E, F, G). This indicates an initial increased rate of differentiation, which depletes the reserve cell pool, which in turn causes a drop in the rate of differentiation.

**Figure 6:**
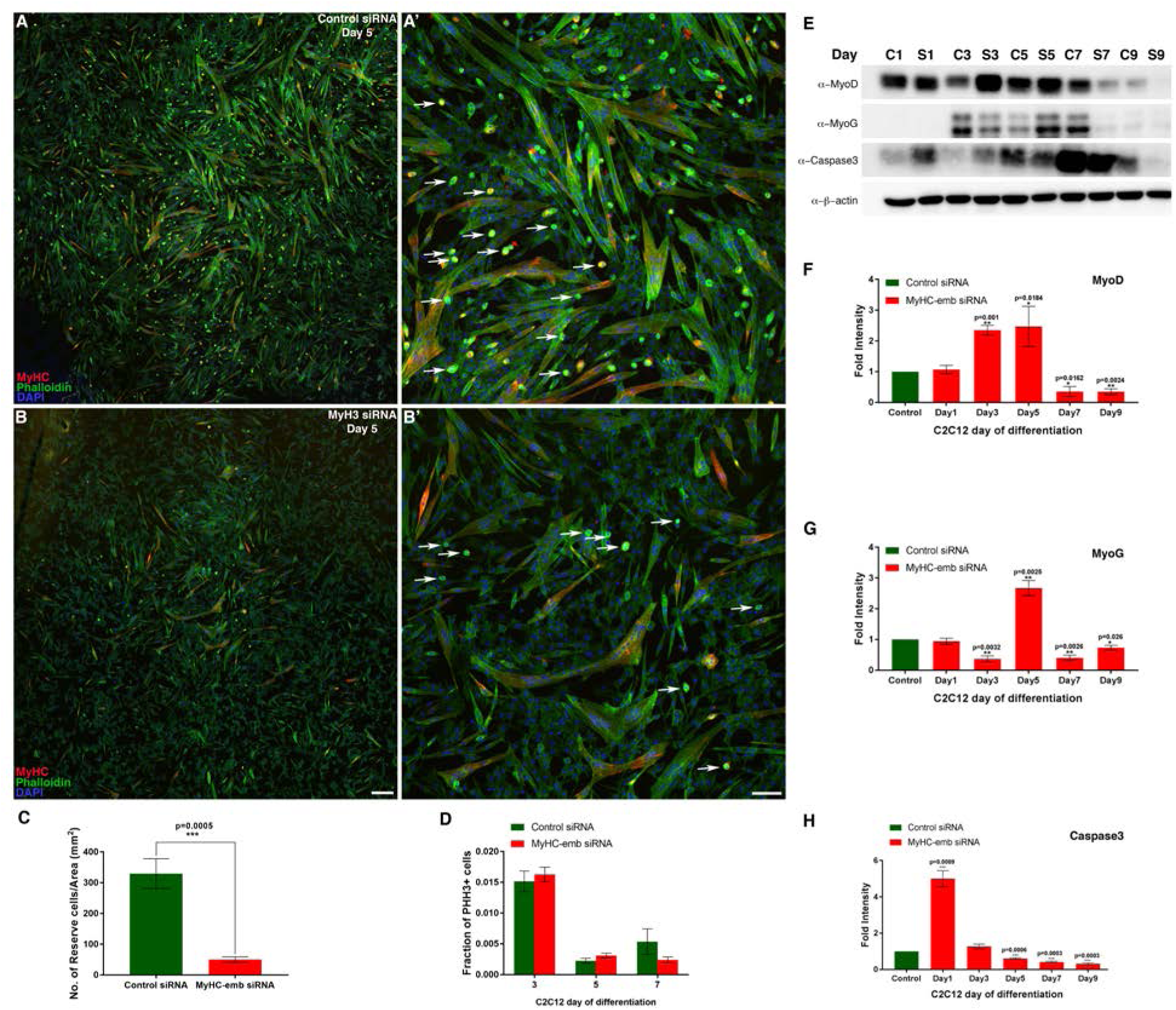
MyHC-embryonic is required for proper myogenic differentiation in vitro. MyHC-emb knockdown in C2C12 cells by siRNA leads to a reduction in the reserve cell population (B, B′) compared to control siRNA treated cells (A, A′), by immunofluorescence for Myosin heavy Chain (Red), Phalloidin (Green), and DAPI (Blue), on day 5 of differentiation, with the white arrows marking some reserve cells in A′ and B′ which are magnified areas taken from A and B respectively. Quantification of the number of reserve cells in control siRNA treated cells compared to *MyH3* siRNA treated cells at day 5 of differentiation indicates that there is a significant, 6-fold decrease in reserve cell number normalized to total area in the *MyH3* siRNA treated cells (C). No significant difference in the fraction of dividing, phospho-histone H3+ cells were observed in the *MyH3* siRNA treated C2C12 cells compared to control siRNA treated cells at days 3, 5 and 7 of differentiation (D). Upon knockdown of MyHC-emb in C2C12 cells, MyoD the protein level increases significantly by 2-fold on early days (days 3 and 5) of C2C12 cell differentiation while the levels decrease significantly to about a quarter of normal levels on later days (days 7 and 9) of differentiation as detected by western blots and densitometry (E and F). The protein levels of Myogenin decreases to a quarter of the normal level on day 3, increases significantly by more than 2-fold by day 5 of C2C12 cell differentiation and again decreases significantly on later days (days 7 and 9) of differentiation as detected by western blots and densitometry upon knockdown ofMyHC-emb (E and G). Cell death as determined by levels of cleaved Caspase3 was significantly elevated in the *MyH3* siRNA treated C2Cl2 cells compared to control siRNA treated cells at day 1 of differentiation, whereas it was significantly less in the *MyH3* siRNA treated cells compared to control cells at days 5, 7 and 9 of differentiation, with no significant difference on day 3 (E), which is quantified by densitometry (H). (Scale bar in B is 100 microns and B′ is 25 microns).

Loss of MyHC-emb during embryonic and fetal myogenesis has striking parallels with the knockdown of MyHC-emb during C2C12 myogenic differentiation, where initially elevated protein levels of MyoD and Myogenin are observed, followed by reduction in their levels (Fig5A, B, C, and Fig6E, F, G). The myogenic progenitor equivalent reserve cells are depleted in C2C12 cells as is seen in vivo during fetal myogenesis, where the Pax7+ myogenic progenitor cells are depleted (Fig6A, A′, B, B′, C and Fig5H, H′, H′′, I, I, I′′′, L).

### Non-cell autonomous signals mediate MyHC-embryonic function in vitro

Next, we wanted to test whether non-cell autonomous signals mediated by MyHC-emb are critical for normal myogenic differentiation. For this, C2C12 cells were treated with conditioned media (from *MyH3* or control siRNA treated C2C12 cells allowed to differentiate for 5 days), differentiated for 4 days, and the fusion index calculated. Intriguingly, we observed that C2C12 cells treated with the *MyH3* siRNA conditioned media had larger and increased number of myofibers compared to control siRNA conditioned media (Fig7B and A respectively). Upon quantification, the *MyH3* siRNA conditioned media treated cells had a statistically significant increase in fusion index as compared to the control siRNA conditioned media treated cells (Fig7C). Thus, culturing C2C12 cells in conditioned media taken from MyHC-emb knockdown cells promoted differentiation, indicating that non-cell autonomous secreted signals mediated by MyHC-emb are critical for regulating myogenic differentiation.

**Figure 7:**
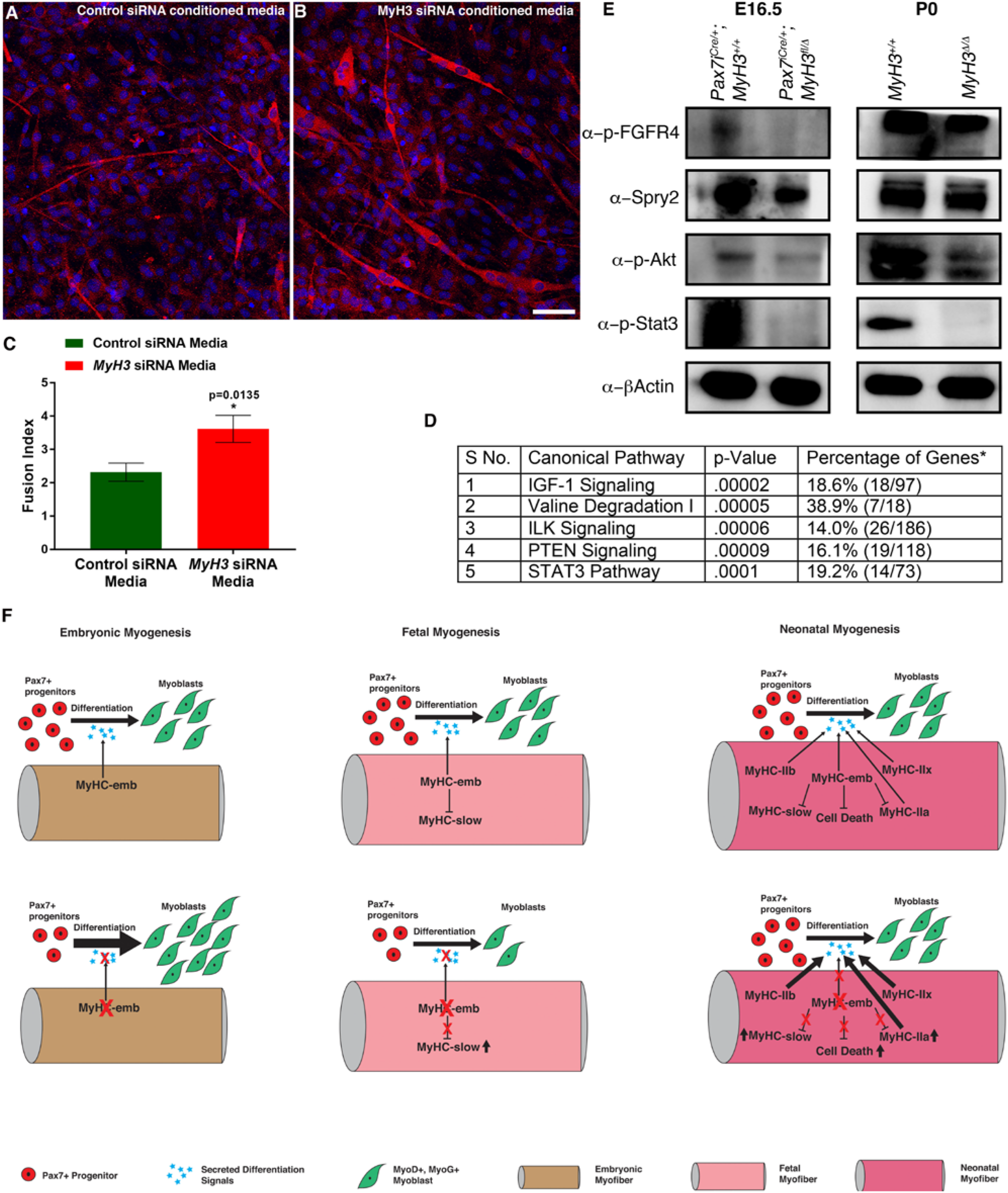
MyHC-embryonic regulates myogenic differentiation non-cell autonomously. C2CI2 cells were allowed to differentiate for 4 days in conditioned media from control siRNA treated cells (A), as compared to *MyH3* siRNA treated cells (B), and labeled by immunofluorescence for MyHC-slow and fast MyHCs to mark the myofibers (red) and nuclei are labeled by DAPI (blue). The cells grown in conditioned media from *MyH3* siRNA treated cells promotes differentiation, with larger myofibers with more nuclei visible (B). The fusion index was calculated and a significant increase in fusion index was observed in the cells differentiated in *MyH3* siRNA treated conditioned media as compared to cells differentiated in conditioned media from control siRNA treated cells (C). The RNA-Seq data from P0 *MyH3^+/+^* and *MyH3^Δ/Δ^* samples for quadriceps, tibialis anterior, gastrocnemius, and diaphragm muscles was subjected to pathway analysis to identify the major pathways altered upon loss of MyHC-emb, where the top 5 pathways that were significantly altered had 4 pathways that are related to growth factor related MAP kinase signaling (D). The p-value and the number of genes affected as compared to total number of genes in the respective pathway are represented in the table (D). In order to validate pathway activation during fetal myogenesis where MyHC-emb plays significant roles, as compared to P0 stage, western blots were performed on E16.5 *Pax7^iCre/+^, MyH3Δ/fl3-7* compared to *Pax7^iCre/+^, MyH3+/+* animals, and *MyH3^Δ/Δ^* and *MyH3^+/+^* animals at P0 for p-FGFR4, Spry2, p-Akt, and p-Stat3 (E). p-FGFR4 levels are drastically downregulated at E16.5 upon loss of MyHC-emb, while it is downregulated upon loss of MyHC-emb at P0 (E). Spry2 levels are downregulated upon loss of MyHC-emb at E16.5, but is unchanged at P0 (E). p-Akt and p-Stat3 levels are downregulated upon loss of MyHC-emb at both E16.5 and P0 stages, with beta-actin used as loading control (E). MyHC-emb thus plays roles in myogenesis during embryonic, fetal and neonatal stages, which are summarized (F). During embryonic myogenesis, MyHC-emb expression in the myofibers facilitates differentiation from progenitors to myoblasts to proceed at a normal rate, mediated by non-cell autonomous, secreted signals from the myofibers (F). In the absence of MyHC-emb during embryonic myogenesis, lack of these non-cell autonomous, secreted signals lead to rapid differentiation of myogenic progenitors to myoblasts, causing depletion of myogenic progenitors and increase in myoblasts (F). MyHC-embc expression in the myofibers during fetal myogenesis is also necessary to facilitate differentiation from progenitors to myoblasts and myoblasts to myofibers, mediated by non-cell autonomous, secreted signals from the myofibers (F). Loss of MyHC-emb function during fetal myogenesis causes depletion of myogenic progenitor and myoblast numbers, presumably due to lack of MyHC-emb mediated secreted signals that allow differentiation to proceed at a normal pace (F). In addition, MyHC-slow protein levels are upregulated at E16.5 upon loss of MyHC-emb (F). During neonatal myogenesis, MyHC-emb levels start to be downregulated and is replaced by the mature fast MyHCs, which together facilitate normal rate of myogenic differentiation by non-cell autonomous, secreted signals (F). MyHC-emb is required to downregulate levels of MyHC-slow and -IIa, as well as reduce cell death (F). In the absence of MyHC-emb during neonatal myogenesis at P0, MyHC-slow and -IIa protein levels are upregulated, and cell death is elevated (F). The increased levels of MyHC-IIa, -slow and possibly other fast MyHCs are able to compensate for the lack of MyHC-emb and no significant effects on myogenic progenitor or myoblast differentiation is observed during this stage (F). (Scale bar in B is 25 microns).

### MAPKinase pathway and FGF signaling are mis-regulated upon loss of MyHC-embryonic

Since MyHC-emb in myofibers non-cell autonomously regulates differentiation of neighboring progenitors, we analyzed our RNA-Seq results, comparing P0 *MyH3^+/+^* and *MyH3^Δ/Δ^* muscles to identify specific signaling pathways that are mis-regulated upon loss of MyHC-emb. We found that of the top 5 pathways that were mis-regulated, 4 were related to MAPkinase signaling (Fig7D). These were the IGF1 (insulin-like growth factor1), ILK (integrin like kinase), PTEN (phosphatase and tensin homolog), and STAT3 (signal transducer and activator of transcription3) pathways respectively (Fig7D).

One of the signaling pathways that mediate differentiation and maintenance of stem cell pool is the FGF (fibroblast growth factor) pathway, where signaling is mediated through MAPkinase activation (Pawlikowski, et al., 2017; Goetz and Mohammadi, 2013; Tsang and Dawid, 2004). FGFR4, a FGF family receptor is central to myogenic differentiation during development, playing a key role in regulating the rate of differentiation of myogenic progenitors and myoblasts (Lagha, et al., 2008; Marics, et al., 2002). Since embryonic or fetal loss of MyHC-emb led to altered rate of myogenic differentiation and several MAPK related pathways were misregulated upon loss of MyHC-emb in neonatal muscles, we hypothesized that signaling mediated by FGFR4 or related receptors might be involved (Lagha, et al., 2008; Marics, et al., 2002). To test this, we carried out western blots for activated forms of known FGF pathway members and found that during fetal myogenesis (E16.5), p-FGFR4, p-Akt and p-Stat3 levels were decreased upon loss of MyHC-emb in the myogenic lineage, indicating that the FGF pathway is mis-regulated upon loss of MyHC-emb during fetal myogenesis (Fig7E). We observed a similar decrease in Spry2, a target of FGF signaling at E16.5 (Fig7E). Interestingly, by P0, p-FGFR4 and Spry2 levels were similar to control levels, while p-Akt and p-Stat3 levels continued to be reduced upon loss of MyHC-emb (Fig7E). This most likely reflects a compensatory effect of adult fast MyHCs, which are expressed by P0, partially rescuing the effect of loss of MyHC-emb by neonatal stages. Thus, loss of MyHC-emb during fetal myogenesis in the myofibers leads to mis-regulation of signaling intermediates in the FGF pathway, which is partially rescued by neonatal stages, which could be the pathway regulating altered rate of differentiation of myogenic progenitors and myoblasts during development.

## Discussion

MyHC-emb is one of three MyHCs expressed in the skeletal muscle during mammalian development, although very little is known about the function of MyHC-emb during various phases of myogenesis during development. Using a mouse conditional targeted allele for MyHC-emb and in vitro tools, we show that MyHC-emb plays novel cell autonomous and non-cell autonomous roles during myogenesis in vivo and in vitro.

### MyHC-embryonic is expressed at high levels during early stages of myogenic differentiation

MyHC-emb and -slow have been reported to be the earliest MyHCs expressed during developmental myogenesis (Lyons, et al., 1990). We found that although both MyHC-emb and - slow transcript expression peaked at E15.5 in vivo and day 6-7 of differentiation in vitro, MyHC-emb was expressed several folds higher compared to MyHC-slow. At the protein level, MyHC-emb exhibited two peaks of expression, at E14.5 and E16.5, whereas MyHC-slow expression was relatively stable until E17.5, when it peaked. Since the peak of transcript expression did not always correlate with protein levels, it is likely that in addition to direct transcriptional regulation of MyHC-emb and -slow, post-transcriptional, translational, or post-translational regulatory mechanisms might also be involved (Schiaffino and Reggiani, 1996; Cox, et al., 1991; Lyons, et al., 1990; Lawrence, et al., 1989; Vivarelli, et al., 1988). While MyHC-emb and -slow proteins label all myofibers during embryonic myogenesis (E13.5), MyHC-slow is restricted to a subset of MyHC-emb+ fibers during fetal myogenesis (E16.5), agreeing with previous reports on primary and secondary myofibers and how fiber type is specified during development (Schiaffino and Reggiani, 1996; DeNardi, et al., 1993; Gunning and Hardeman, 1991). Presumably, fetal myofibers that express MyHC-emb and -slow give rise to adult MyHC-slow+ fibers, while those fetal fibers that express MyHC-emb and MyHC-peri (which we did not investigate) give rise to adult fast fibers (Schiaffino and Reggiani, 1996; DeNardi, et al., 1993; Gunning and Hardeman, 1991). Thus, functionally MyHC-emb might have significant roles in all myofibers during embryonic and fetal myogenic phases, while MyHC-slow might be important in all myofibers during embryonic phase and only in MyHC-slow+ fibers during fetal myogenesis.

### Loss of MyHC-embryonic has different effects on different muscles

One of our most striking observations was that although MyHC-emb is expressed in all muscles during the course of development, the effect of loss of MyHC-emb was different in different muscles. We found this to be the case when we studied the effect of loss of MyHC-emb on mis-regulation of other MyHCs, fiber number, size and type, and the candidate gene transcripts that are mis-regulated upon loss of MyHC-emb. Each of these are discussed further in the 3 sections below.

Previous studies on MyHC-IIb and -IIx knockouts suggest that other MyHCs compensate for loss of a specific MyHC in adult stages (Allen and Leinwand, 2001; Sartorius, et al., 1998). Although there were no major changes in the temporal or spatial expression patterns of other MyHCs upon loss of MyHC-IIb or IIx, an increase in MyHC-IIx upon loss of MyHC-IIb and MyHC-IIa upon loss of MyHC-IIx were reported, which might indicate that inbuilt compensatory mechanisms exist to express MyHCs with closest contractile properties upon loss of function of a specific MyHC (Allen and Leinwand, 2001; Sartorius, et al., 1998). Another intriguing possibility is that the MyHC that lies closest in the MyHC gene cluster to the MyHC that is deleted takes over its function. We investigated whether such compensatory upregulation occurs during developmental myogenesis upon loss of MyHC-emb and found that transcripts of MyHC-IIa, the MyHC gene closest to MyHC-emb in the gene cluster is upregulated in all muscles tested neonatally, MyHC-IIx which is the second closest gene and MyHC-peri which is the fourth gene in the cluster are upregulated in 2 muscles each. This raises two interesting possibilities-regulatory elements might be shared between MyHC-emb and the MyHCs neighboring it, or a compensatory upregulation similar to that seen in the case of Hox clusters with respect to limb colinearity might be involved (Kmita and Duboule, 2003). Transcript levels of MyHC-slow, which is not part of the fast MyHC gene cluster, is reduced in 2 muscles at neonatal stages upon loss of MyHC-emb, possibly due to a decrease in the total number of fibers in those muscles. Such extensive mis-regulation of other MyHCs was not observed in MyHC-IIb or -IIx null mice, indicating that MyHC-emb, as a developmental MyHC is crucial in establishing normal levels of both fast and slow MyHCs during neonatal stages. In addition, although MyHC-emb is expressed in all muscle fibers, loss of MyHC-emb does not lead to a uniform effect on other MyHCs in all muscles (except for MyHC-IIa, which was upregulated in all muscles studied). This is most likely indicative of the fiber type diversity between specific muscles.

Loss of MyHC-emb led to increased number of myofibers in the fast fiber-rich EDL muscle, with no effect on the slow fiber-rich soleus muscle at neonatal stages. On the other hand, the proportion of myofibers with the least cross-sectional area were increased upon loss of MyHC-emb in the soleus, without having any effect on the EDL. Thus, MyHC-emb regulates myofiber number in fast muscles and myofiber area in slow muscles. Previously it was shown in MyHC-IIb null mice that the mean fiber size was increased significantly with fibers appearing of uniform size, and in MyHC-IIx null mice where the proportion of very small and very large fibers were significantly increased (Acakpo-Satchivi, et al., 1997). Further, specific adult muscles were affected to different degrees with respect to fiber size and interstitial fibrosis, for instance the gastrocnemius and quadriceps were most affected in MyHC-IIb null mice whereas the TA and EDL were most affected in IIx null mice, indicating that different MyHC isoforms play distinct roles (Allen, et al., 2001; Allen, et al., 2000). We also found that fusion index, a measure of efficiency of myogenic differentiation, was significantly reduced upon knockdown of MyHC-emb in vitro, supporting our in vivo findings. Most interesting was the observation that the number of MyHC-slow+ fibers and the total levels of MyHC-slow protein were drastically increased in MyHC-emb null neonatal muscles. At the transcript level, we found a significant reduction in MyHC-slow levels whereas at the protein level, we saw that levels increased upon loss of MyHC-emb, both in vivo and in vitro. This inverse correlation suggests that MyHC-slow is not simply regulated at the transcriptional level, but other post-transcriptional, translational, or post-translational mechanisms might also play a role. Previous studies have reported an inverse correlation between MyHC transcript and protein levels, although the precise mechanism has not been investigated and could be related to changes in mRNA or protein stability, or rate of translation (Schiaffino and Reggiani, 1996; Cox, et al., 1991; Lyons, et al., 1990; Vivarelli, et al., 1988). The decrease in fiber number and increased proportion of smaller fibers that are observed neonatally in specific muscles upon loss of MyHC-emb could be a direct consequence of the reduced number of myogenic progenitors and myoblasts upon fetal loss of MyHC-emb function (discussed in the next section). However, this effect is not uniform across all muscles. One reason for this could be to do with neonatal compensation by specific adult MyHC isoforms, which start being expressed at neonatal stages. Since MyHC-emb is expressed in both future fast and slow fibers, it is plausible that it has effects on muscles rich in both kinds of fibers. How it has distinct effects on these two types of muscles is unclear, and might have to do with expression levels of the specific MyHCs that are compensating for the loss of MyHC-emb in that specific muscle, which is something that needs further investigation.

An RNA-Seq experiment to identify target genes that are mis-regulated upon loss of MyHC-emb in identified candidates broadly related to general myogenic differentiation or muscle structure. 12 candidates (8 shared between different muscles and 4 unique to a specific muscle) that we identified were also discovered in a previous study to uncover transcriptional regulators involved in early muscle differentiation using C2C12 mouse myoblasts (Rajan, et al., 2012). These results substantiate that MyHC-emb is essential for normal myogenic differentiation and loss of MyHC-emb leads to mis-regulation of genes related to myogenic differentiation. Further, 162 genes were uniquely mis-regulated in any of the 4 muscles, validating that loss of MyHC-emb has distinctive effects in different muscles, reflecting variations in fiber type between the different muscles.

### MyHC-embryonic is a key regulator of embryonic, and fetal myogenesis

We found that MyHC-emb is crucial for proper myogenic differentiation in both embryonic and fetal phases of developmental myogenesis. However, there were differences observed upon loss of MyHC-emb between these phases, which possibly is a reflection of the differences in myogenic progenitor and myoblast behavior between embryonic and fetal phases of myogenesis. During embryonic myogenesis at E13.5, we found that loss of MyHC-emb led to a decrease in Pax7 (myogenic progenitor) protein levels and an increase in MyoD and Myogenin (myoblast) protein levels, indicating that MyHC-emb increases the rate of myogenic progenitor to myoblast differentiation at this phase. At E16.5, during fetal myogenesis, loss of MyHC-emb led to a decrease in protein levels of both Pax7 (myogenic progenitor) and MyoD and Myogenin (myoblast), indicating MyHC-emb increases the rate of differentiation of myogenic progenitors to myoblasts and myoblasts to myofibers. We validated this by quantifying the number of myogenic progenitors and myoblasts at E16.5, where we saw a striking reduction in the number of both of these cell types upon loss of MyHC-emb. None of this was due to elevated cell death or altered rate of cell division. We also confirmed that MyHC-emb is not expressed by myogenic progenitors or myoblasts and is only expressed by myofibers, implying that the observed effect on differentiation has to be non-cell autonomous, where signals from the myofibers lacking MyHC-emb lead to rapid rate of differentiation of the myogenic progenitors, depleting the myogenic progenitor pool. Interestingly, a recent study reported that the Pax7+ muscle stem cell pool and its differentiation is regulated by signals from the differentiated myofibers, including mechanical forces of muscle contraction (Esteves de Lima, et al., 2016).

Although the myogenic progenitor pool depletion upon loss of MyHC-emb was consistent across embryonic and fetal phases of myogenesis, the myoblast pool behaved differently, presenting elevated levels during embryonic myogenesis and reduced levels during fetal myogenesis. The most likely explanation for this is the inherent differences between embryonic and fetal myogenic progenitors, which have been previously described (Biressi, et al., 2007a; Biressi, et al., 2007b; Stockdale, 1992). Briefly, embryonic myogenic progenitors start off differentiating in an environment where there are no myofibers, are more prone to differentiation in culture, and are not inhibited by secreted signals such as Transforming Growth Factor-β (TGF-β) and Bone Morphogenetic Protein-4 (BMP-4) in culture, compared to fetal progenitors (Murphy and Kardon, 2011; Biressi, et al., 2007a). Also, during embryonic myogenesis, a small subset of embryonic myogenic progenitors differentiate into primary myofibers and the rest are kept in a committed state for differentiation into secondary myofibers during fetal myogenesis (Biressi, et al., 2007a). Thus, it is possible that the low number of progenitors and myoblasts that are differentiating during embryonic myogenesis ensure that levels of the myoblast progenitors increase during this phase. However, during fetal myogenesis, both the progenitor and myoblast populations are differentiating rapidly, which leads to depletion of both populations of cells. Further, myofibers from embryonic myogenesis are thought to be “slow” type and from fetal myogenesis are thought to be “fast” type (Biressi, et al., 2007a). Our results indicating that both MyHC-slow and fast (MyHC-IIa, -IIx and -peri) are upregulated neonatally suggests that both embryonic and fetal myogenesis are affected upon loss of MyHC-emb. In addition, C2C12 cells have been proposed to behave similarly to fetal myoblasts (Biressi, et al., 2007b). MyHC-emb depletion in C2C12 myogenic cells led to a reduction of myogenic progenitor equivalent reserve cell pool and decrease in protein levels of myoblast markers MyoD and Myogenin at later time points of differentiation, corroborating our in vivo findings.

This effect on the rate of differentiation of myogenic progenitors and myoblasts was not observed at P0, suggesting that other MyHCs compensate for loss of MyHC-emb by this stage, fitting well with our observation that MyHC-IIa, and -slow protein levels, and MyHC-IIx and -perinatal transcript levels are elevated by this stage.

### MyHC-embryonic functions through non-cell autonomous signals mediated by the FGF-MAPKinase pathway

Both positive and negative signals have been proposed to regulate the rate of differentiation of myogenic progenitors and myoblasts (Bryson-Richardson and Currie, 2008). We and others have previously shown that extrinsic signals from the muscle connective tissue fibroblasts regulate muscle differentiation and fiber type (Nassari, et al., 2017; Mathew, et al., 2011; Murphy, et al., 2011; Hasson, et al., 2010; Joe, et al., 2010).

Using conditioned media from C2C12 cells treated with *MyH3* siRNA, we validated that secreted factors from the myofibers are involved in non-cell autonomous regulation of myoblasts upon loss of MyHC-emb. A pathway analysis of the RNA-Seq data comparing neonatal MyHC-emb null muscles to control muscles indicated that MAPKinase pathways were dysregulated upon loss of MyHC-emb. Among numerous signaling pathways where MAPKinase signaling is involved, we found that phenotypes described with respect to signaling mediated by the Fibroblast Growth Factor Receptor-4 (FGFR4), bear a striking resemblance to the myogenic differentiation defects observed by us (Lagha, et al., 2008; Marics, et al., 2002).

Numerous examples indicate that the FGF signaling pathway is crucial for proper myogenesis. FGFs have been reported to downregulate myogenin levels to regulate myogenic differentiation (Fox and Swain, 1993). In zebrafish, FGF8 signaling regulates Pax3, Pax7 and MyoD expression as well as terminal differentiation of a subset of somitic muscles (Hammond, et al., 2007; Groves, et al., 2005). In the head muscles in chick, FGF8 regulates the choice between proliferation and differentiation of muscle precursors (von Scheven, et al., 2006). FGFR4 is expressed at high levels in dividing Pax3/Pax7+ myogenic progenitors and myoblasts in the chick and inhibition of FGFR4 led to down-regulation of MRFs such as Myf5 and MyoD and muscle progenitor differentiation arrest, which could be rescued by exogenous FGF (Ben-Yair and Kalcheim, 2005; Marics, et al., 2002). In FGF6 null mice, the soleus muscle hypertrophied following regeneration possibly mediated by Insulin-like Growth Factor (IGF) signaling, and FGF6 has been suggested to regulate myoblast proliferation versus differentiation based on the specific receptor it interacts with (Armand, et al., 2006; Armand, et al., 2004). Lack of FGFRL, a fibroblast growth factor like receptor which interacts with FGF ligands leads to embryonic loss of slow muscle fibers in mice, while FGF6 knockout adult mice had increased numbers of MyHC-slow+ fibers following regeneration (Amann, et al., 2014; Armand, et al., 2005). FGFR4 is strongly expressed in adult differentiating myoblasts and loss of FGFR4 led to aberrant adult muscle regeneration characterized by delayed and poorly coordinated differentiation, fat infiltration, inflammation and calcification (Zhao, et al., 2006; Zhao and Hoffman, 2004). FGFR4 is also a direct target of Pax3, crucial in regulating the equilibrium between myogenic progenitors and committed myoblasts during differentiation (Lagha, et al., 2008). In Pax3 null embryos where FGFR4 expression is lost in the myogenic lineage, the FGF signaling pathway is mis-regulated, with reduced phospho-Akt, and phospho-P38 levels and elevated phospho-ERK levels (Lagha, et al., 2008). By overexpressing an FGF signaling downstream intermediate-Spry2 in the Pax3 lineage during mouse embryonic development, phospho-ERK levels were downregulated, and even more significantly, an increase in the number of Pax7+ myogenic progenitors was observed as compared to the Myogenin+ committed myogenic cells (Lagha, et al., 2008). Overexpression of FGF4 leads to downregulation of muscle markers including Pax3 and MyoD, loss of FGFR4 expression and inhibition of terminal differentiation in limb muscles during chick development (Edom-Vovard, et al., 2001). FGFR4 has been shown to be mutated and activated in rhabdomyosarcoma, a sarcoma where cells exhibit myoblast-like characteristics, where its target STAT3 was also shown to be activated (Li, et al., 2013; Taylor, et al., 2009). FGF signaling mediated by ERK, P38, PI3Kinase and Stat signaling is known to be involved in satellite cell self-renewal, and skeletal muscle regeneration (Pawlikowski, et al., 2017). Strikingly, missense mutations in human FGFR3 have been reported to cause CATSHL syndrome, where patients exhibit camptodactyly and scoliosis, features which are also observed in Freeman-Sheldon and Sheldon-Hall syndromes caused by mutations in the MyHC-emb encoding *MyH3* gene, indicating that MyHC-emb and FGF signaling might have common functions during development (Toydemir, et al., 2006a; Toydemir, et al., 2006c).

In order to verify whether FGFR4 mediated signaling is involved in regulating the non-cell autonomous myogenic differentiation defects that we observed, we tested the FGFR4 pathway in fetal and neonatal muscle samples lacking MyHC-emb function, as compared to control samples. We found that the p-FGFR4 itself and FGFR4 pathway mediators such as p-Akt, p-Stat3 and Spry2 levels were decreased upon loss of MyHC-emb in the myogenic lineage at fetal stages, strongly suggesting that aberrant FGF signaling leads to the observed differentiation defects during fetal myogenesis. These effects are partially rescued by neonatal stage, possibly by compensatory effects of other MyHCs, which presumably is why the non-cell autonomous differentiation defects are not observed at this stage.

## Model and Conclusions

Based on our findings, we propose that MyHC-emb has critical cell autonomous and non-cell autonomous functions during embryonic, fetal and neonatal stages of myogenesis. During embryonic myogenesis, MyHC-emb is required to non-cell autonomously regulate the rate of differentiation of myogenic progenitors to myoblasts. In the absence of MyHC-emb during embryonic myogenesis, signals from the myofibers cause the myogenic progenitors to differentiate rapidly, leading to depletion of the progenitor pool and increase in the myoblast pool (Figure7F).

During fetal myogenesis, MyHC-emb non-cell autonomously regulates the rate of differentiation of myogenic progenitors to myoblasts and myoblasts to myofibers. MyHC-emb is required to cell autonomously regulate levels of MyHC-slow during fetal myogenesis. Thus, in the absence of MyHC-emb during fetal myogenesis, there is elevated levels of MyHC-slow and signals from the myofibers cause the myogenic progenitors and myoblasts to differentiate rapidly, leading to depletion of both cell pools (Figure7F). During neonatal myogenesis, MyHC-emb has cell autonomous effects, and possibly also non-cell autonomous effects in regulating the rate of differentiation. It is likely that mature MyHCs that begin expressing at neonatal stages compensate for MyHC-emb which could explain why non-cell autonomous effects of MyHC-emb are not observed at neonatal stage. Within the myofibers, MyHC-emb is required to regulate myofiber number in fast muscles, myofiber area in slow muscles, number of MyHC-slow+ fibers, levels of MyHC-slow, -IIa, -IIx and -perinatal, and cell death (Figure7F). Thus, MyHC-emb is a critical regulator of all phases of developmental myogenesis, performing distinct and overlapping functions at embryonic, fetal and neonatal myogenesis.

Whether the secreted signals mediated by MyHC-emb, that non-cell autonomously regulate myogenic differentiation is the same across embryonic, fetal and neonatal myogenesis is an open question. However, based on previous results and our data, MAPKinase signaling downstream of the FGF pathway, specifically FGFR4 receptor-mediated signaling, is at least partially responsible for the observed effects (Lagha, et al., 2008; Marics, et al., 2002). It is also likely that positive and negative signals are involved in this and a balance between the two is required to facilitate normal pace of myogenic differentiation. In the absence of MyHC-emb, this equilibrium is shifted to increase the rate of differentiation and depletion of the corresponding pools of myogenic cells. Although not straightforward, it will be fascinating to identify all possible signals that mediate the non-cell autonomous effect of MyHC-emb on myogenic progenitors and myoblasts, employing in vitro and in vivo approaches. This should not only shed light on developmental myogenesis but also lead to a better understanding of the congenital contracture syndromes associated with mutations in MyHC-emb.

## Materials and Methods

### Mice

*MyH3^fl3-7/+^* mice were generated by flanking exons 3-7 of MyHC-emb with LoxP sites, according to published protocols (Wu, et al., 2008). Briefly, an 11.3 Kb genomic fragment of mouse MyHC-emb was recombineered into the pStartK vector from BAC clone RP23-67L23 (Children’s Hospital Oakland Research Institute - CHORI). By a series of recombineering and cloning steps, LoxP sites, FRT-PGKNeo-FRT and restriction enzyme sites were added to the construct, and validated (Wu, et al., 2008). The targeting construct was transferred to the pWS-TK2 vector, linearized and used for integration into ES cells following positive and negative selection (Wu, et al., 2008). Genomic DNA from about 200 ES cell clones were screened for 5′ and 3′ LoxP integration by PCR, and 24 clones were identified that were positive for both 5′ and 3′ targeting events. From these, 2 clones were chosen for microinjection into blastocysts, chimeric animals from these crossed to C57Bl/6J wild type animals and the offspring genotyped for targeting. Once targeted animals were identified, they were bred for 5-6 generations to bring them into the C57Bl/6J background, and the *neo* cassette removed by crossing to the *R26R^Flpe^* mice (Farley, et al., 2000). *MyH3^Δ/+^* mice were generated by crossing the *MyH3^fl3-7/+^* mice to the ubiquitous Cre-expressing *Hprt^Cre^* mice (Tang, et al., 2002). The *MyH3^fl3-7/+^* mice were generated at the transgenic core, University of Utah, Salt Lake City, UT, USA.

Other Cre-drivers used were *Pax3^Cre^* (Engleka, et al., 2005), and *Pax7^iCre^* (Keller, et al., 2004). C57Bl/6J wild type mice were used in this study. All of the animal maintenance and experiments were performed according to Institutional Animal Care and Use Committee (IACUC) approved protocols of the University of Utah. Genotyping was carried out by PCR using genomic DNA extracts prepared from mouse ear clips, and primer sequences as well as PCR protocols used for genotyping can be provided upon request.

### Cell culture

C2C12 mouse myoblasts (ATCC; Cat# CRL-1722) were cultured and maintained according to ATCC guidelines in growth medium containing DMEM-Dulbecco’s Modified Eagle Medium (Gibco; Cat# 11995065) supplemented with 10% (v/v) fetal bovine serum (Sigma; Cat# F2442) and 2% penicillin-streptomycin (Gibco; Cat# 15140122). C2C12 cells were differentiated in differentiation media containing DMEM, 2% (v/v) horse serum (BioAbChem; Cat# 72-0460) and 2% penicillin-streptomycin.

For immunofluorescence studies, about 30,000 C2C12 myoblasts were seeded onto gelatin-coated coverslips (Neuvitro; Cat# GG-12) in 24-well plates (Nunc; Cat# 142485) in growth media, which was replaced with differentiation media after 24 hours to facilitate differentiation. Subsequently, cells were maintained in differentiation medium; 50% medium replacement was performed daily in all the wells and cells were harvested from 3 wells in triplicates each day until the end of the assay. These experiments were carried out a minimum of three times. For RNA isolation and protein lysate preparation from C2C12 cells over the course of differentiation, the same protocol was used, except that the cells were seeded directly in the 24-well plate without the gelatin-coated coverslip.

For knocking down MyHC-emb expression in C2C12 myoblasts during myogenic differentiation, reverse transfection was performed. Briefly, 30,000 C2C12 cells were seeded in each well of a 24-well plate (onto gelatin coated coverslips in the well for immunofluorescence) layered with the transfection mix. The transfection mix comprised 100μl Opti-MEM (Gibco; Cat# 31985070), 50nM of *MyH3* or control siRNA (Ambion; Cat# s70258 and 4390847, respectively) and 2μl Lipofectamine RNAiMAX (Invitrogen; Cat# 13778150). The cells were cultured in growth medium for 48 hours to allow cells to grow to ∼80% confluence and efficient transfection. Subsequently, differentiation was induced as explained above. All experiments were performed in triplicates. For fusion index analysis of siRNA treated cells, the coverslips containing cells were harvested on days 3, 5 and 7 of myogenic differentiation. All treatments were carried out uniformly between control and transfected cells.

For the conditioned media experiments, C2C12 cells were cultured in a 24 well dish after treatment with control and *MyH3* siRNA respectively. The conditioned media from these control and *MyH3* siRNA treated cells were collected at day 5 of plating and stored at −20^0^C. Next, ∼30,000 C2C12 cells were plated on gelatin coated coverslips in normal growth media. After 48 hours of growth in normal growth media, the media was replaced with the conditioned media from control siRNA and *MyH3* siRNA treated cells respectively, and allowed to grow for another 4 days, following which the coverslips were processed for staining, images captured and fusion index calculated.

### RNA isolation, cDNA synthesis and quantitative PCR (qPCR)

RNA was isolated from tibialis anterior, quadriceps, gastrocnemius and diaphragm muscles of *MyH3^+/+^* and *MyH3^Δ/Δ^* mice at post-natal day 0 (P0) using RNeasy Lipid Tissue Mini Kit (Qiagen; Cat# 74804). RNA was also isolated from C2C12 myoblasts, harvested on specific days of differentiation as described in the cell culture section above, using RNeasy mini kit (Qiagen; Cat# 74106). cDNA was prepared using 1μg of RNA using SuperScript III reverse transcriptase (Invitrogen, Cat# 18080-044) and oligo (dT) (Invitrogen; Cat# 58862) according to manufacturer’s recommended protocol. qPCR was performed using SYBR Green with ROX as internal reference (Applied Biosystems; Cat# 4367659) using the ABI 7500 Fast Real Time PCR system (Applied Biosystems). The genes studied and primers used for gene expression quantification are listed in Supplementary Table 1. All the qPCR assays were performed as recommended by ABI using 100nM of forward and reverse primers of the respective genes in a final reaction volume of 15μl. Each reaction was performed in triplicates and normalized to *Gapdh* transcript levels. The expression of target genes in the mutant muscles and *MyH3* siRNA transfected cells were normalized to that of wild type muscles and control siRNA transfected cells respectively (Livak and Schmittgen, 2001). Three technical replicates were taken for the gene expression analysis. A minimum of four biological replicates of mutant and knock out mouse muscle samples and three biological replicates of *MyH3*and control siRNA transfected samples were used for performing the expression analysis. The error bar represents the standard error of the mean.

### RNA-Seq

Quadriceps, tibialis anterior, gastrocnemius, and diaphragm muscles from 6 *MyH3^+/+^* and *MyH3^Δ/Δ^* animals respectively were harvested at post-natal day 0 (P0) and RNA extracted using RNeasy Lipid Tissue Mini Kit according to manufacturers protocols (Qiagen; Cat# 74804). Integrity and concentration of the isolated RNA was verified using the 2200 TapeStation (Agilent). Library preparation was performed using the Illumina TruSeq Stranded mRNA sample preparation kit with oligo dT selection according to manufacturer’s protocol and single-end 50 bp reads were generated using an Illumina HiSeq 2000 instrument. Transcript annotations for mm10 (M_musculus, Dec 2011) were used from Ensembl. Reads were aligned using Pysano and annotated splice junctions generated using USeq (www.sourceforge.net/projects/USeq). Splice junction reads were mapped to genomic coordinates using the SamTranscriptomeParser application in USeq. Differential gene expression was identified using the Defined Region Differential Seq (DRDS) application in USeq, following which paired-sample differential gene expression analysis was performed using DESeq2 (Love, et al., 2014). RNASeq metrics were generated using Picard’s CollectRnaSeqMetrics, samples clustered using custom R scripts (Analysis/Plots) and significant genes were run in IPA to generate pathway analyses. Differentially expressed genes identified by RNA-Seq, common between muscles or unique to specific muscles in *MyH3^Δ/Δ^* knockout animals compared to *MyH3^+/+^* animals have been represented by a Venn diagram.

### Western blots

For protein isolation from cell culture samples, C2C12 cells were plated and cultured as described above. Cells were lysed in ice-cold radioimmunoprecipitation assay (RIPA) buffer (Sigma; Cat# R0278-500ml) containing protease inhibitor (Sigma; Cat# P8340-5ml) at 1:100 dilution, at specific time points as indicated. Protein was also isolated from limbs of E13.5 *Pax3^Cre/+^, MyH3+/+*, and *Pax3^Cre/+^, MyH3Δ/fl3-7* embryos, and E16.5 *Pax7^iCre/+^, MyH3+/+*, and *Pax7^iCre/+^, MyH3Δ/fl3-7* embryo hind limbs using Qproteome FFPE Tissue kit (Qiagen; Cat# 37623). Total protein was isolated from the hind limb muscles of *MyH3^+/+^* and *MyH3^Δ/Δ^* neonatal mice at P0 by homogenization using the Precellys 24 homogenizer (Bertin Technologies). Quantification of the protein samples was done using Pierce BCA Protein Assay kit (Thermo Scientific; Cat# 23225) as per manufacturer’s protocol. Protein samples were separated on a 10% SDS-PAGE and the separated proteins were transferred onto a PVDF membrane (Millipore; Cat# iPVH00010) at 4^0^C for 2 hours. Western blot was performed using standard procedures, with blocking in 5% skimmed milk (Himedia; Cat# RM1254-500GM) for 3 hours, washes in PBS containing 0.1% Tween 20 (Sigma; Cat# P7949-100ML), incubated overnight in primary antibody at 4^0^C, and 2 hours in secondary antibody at room temperature. HRP conjugated secondary antibodies were used, and signal was detected using the HRP substrate (Millipore; Cat# WBLUF0100). Blots were imaged using ImageQuant LAS 4000, from GE. Blots were stripped using standard procedures and re-probed for beta-actin. Densitometry was performed to quantify the amount of protein normalized to beta-actin levels by using ImageQuant software, and data from a minimum of 3 independent replicates were used for analysis, with the error bar representing the standard error of the mean. Antibodies used are listed in Supplementary Table 2.

### Immunofluorescence and microscopy

For the fluorescence intensity measurements of MyHC-emb and MyHC-slow, wild type embryos were harvested from timed matings, fixed, and embedded in Optimal Cutting Temperature (OCT) (Tissue-Tek). Neonatal hind limbs were embedded in OCT and flash frozen in 2-methyl butane cooled in liquid nitrogen. Similarly, for knockout or conditional knockout experiments, embryos at the correct stages were harvested from timed matings (E13.5 *Pax3^Cre/+^, MyH3+/+*, and *Pax3^Cre/+^, MyH3Δ/fl3-7* embryos, and E16.5 *Pax7^iCre/+^, MyH3+/+*, and *Pax7^iCre/+^, MyH3Δ/fl3-7*) or hind limbs were flash frozen from *MyH3^Δ/Δ^* and *MyH3^+/+^* neonates at P0. These samples were sectioned at a thickness of 10μm using a cryomicrotome (Thermo Scientific; Microm HM 550) and adjacent sections collected on coated glass slides (VWR; Cat # VWRU48311-703). The sections were processed for immunofluorescence as described below and adjacent sections were used for immunofluorescent based detection of MyHC-emb, MyHC-slow, Pax7, MyoD and Phospho-histone H3. C2C12 cells grown on coverslips, described in cell culture section above, were also processed for immunofluorescent based detection. The tissue sections and C2C12 cells were fixed in 4% paraformaldehyde (PFA) for 20 minutes and washed with PBS. For tissue sections, the antigen was retrieved wherever required (specified in Supplementary Table 2) by heating the slides to 120°C for 5 minutes in citrate buffer (1.8mM citric acid and 8.2 mM sodium citrate in water) in a 2100 PickCell Retriever (Aptum Biologics Ltd). Tissue sections and cells were blocked with 5% goat serum (BioAbChem; Cat# 72-0480) in PBS containing 0.1% Triton-X-100 (MP Biochemicals; Cat# 194854) for 1 hour at room temperature (RT), incubated overnight at 4°C with appropriate concentration of primary antibody listed in Supplementary Table 2, washed thrice with PBS, incubated with secondary antibody for 2 hours at RT, and washed with PBS. Where amplification was required, the samples were incubated with a biotin conjugated secondary antibody for 2 hours at RT and then with Strep coupled fluorophore for 1 hour at RT, and washed thrice with PBS. Samples were post-fixed in 4% PFA, rinsed in distilled water and mounted using DAPI Fluoromount-G (Southern Biotech; Cat# 0100-20). Antibodies used are listed in Supplementary Table 2. Fluorescence microscopy was performed using a Leica TCS SP5 II, or Nikon A1R confocal microscope.

### Cell counts, fiber counts and statistics

For fusion index analysis, randomly selected 5 non-overlapping fields of view (2×3 tiles) per coverslip for control and *MyH3* siRNA transfected cells or cells treated with conditioned media from control and *MyH3* siRNA transfected cells were imaged using identical settings on a Leica TCS SP5 II. Myotubes were labeled for Myosin Heavy Chain using a mixture of My32 and MyHC-slow antibodies and nucleus by DAPI. Counts for total nuclei, number of myotubes and nuclei within the myotubes per unit area were performed using ImageJ (Schindelin, et al., 2012; Schneider, et al., 2012). The fusion index was calculated as the percentage of nuclei within myotubes compared to the total number of nuclei. For the reserve cell counts, *MyH3* and control siRNA treated cells were cultured, stained for Phalloidin, Myosin Heavy Chain (mixture of My32 and MyHC-slow antibodies) and nucleus by DAPI and images captured. Total number of DAPI+ nuclei and Phalloidin+ reserve cells were counted using spot and annotation functions in Imaris software respectively, and normalized to unit area (http://www.bitplane.com/). Phospho-histone H3+ nuclei were counted manually, and the total DAPI+ nuclei were counted using the particle analyzer function in ImageJ (Schindelin, et al., 2012; Schneider, et al., 2012). The fraction of Phospho-histone H3+ nuclei compared to total nuclei is represented. All of these experiments were performed a minimum of 3 times and images captured for both control and *MyH3* siRNA treated cells using identical settings.

The fluorescence intensity for MyHC-emb and MyHC-slow in wild type embryos during development was calculated as follows. Hind limbs from a minimum of 3 embryos per developmental stage (E12.5, E13.5, E14.5, E15.5, E16.5, E17.5 and E18.5) were sectioned and processed for immunofluorescence as described and adjacent sections labeled for MyHC-emb and MyHC-slow. Imaging was performed using identical settings and fluorescence intensity was calculated using ImageJ (McCloy, et al., 2014; Burgess, et al., 2010). The average fluorescence intensity for each developmental stage was calculated across biological replicates, and relative fluorescence intensity was calculated by subtracting the average fluorescence intensity of each stage from E13.5-18.5 from that of E12.5 (McCloy, et al., 2014; Burgess, et al., 2010).

Pax7+ cell counts from the entire P0 hind limb cross section was performed using the spot function in Imaris software (http://www.bitplane.com/). Total P0 hind limb muscle cross sectional area was quantified using the surface function in Imaris software. A minimum of 3 replicates for each genotype were analyzed in this manner with the data represented as mean ± standard error of the mean. MyHC-slow fiber count and fiber area were measured using the Semi-automatic muscle analysis using muscle histology software (SMASH) (Smith and Barton, 2014). MyHC-slow fiber numbers were counted using fiber typing function and muscle fiber area was measured using the fiber properties function of SMASH software respectively in EDL and soleus muscles of control and *MyH3* null mice.

Pax7+, MyoD+ and Phospho-histone H3+ cell counts from the entire E16.5 hind limb cross section was performed using the spot function in Imaris software (http://www.bitplane.com/). Total E16.5 hind limb muscle cross sectional area was quantified using the surface function in Imaris software and the data is presented as number of Pax7+, MyoD+, or Phospho-histone H3+ cells per unit area.

A minimum of 3 replicates for each genotype were analyzed in this manner with the data represented as mean ± standard error of the mean.

Data from all the experiments were analyzed using parametric, unpaired t-test using the Graphpad Prism software. The error in all of the graphs are represented as mean ± standard error of the mean. The p-value is indicated on the graph along with asterisks and p-value ≤ 0.05 is considered significant.

## Author Contributions

AS, MA, AK, PK, MS and SJM carried out the experiments, AS, MA, AK, GK and SJM were involved in experimental design, and SJM wrote the manuscript.

## Acknowledgements

This work was funded by an Intermediate Fellowship (IA/I/13/1/500872) from the Wellcome Trust/DBT India Alliance to SJM and NIH R01HD053728 to GK. We also acknowledge funding from the Department of Biotechnology, and the Regional Centre for Biotechnology (RCB). AS and PK are funded by senior research fellowships from CSIR, MA by senior research fellowship from ICMR, and MS by a Young Investigator award from RCB. We acknowledge valuable suggestions and help from Dr. Kirk Thomas, Dr. Sen Wu, and Dr. Charles Murtaugh at the University of Utah with the *MyH3* targeting design. *MyH3* targeting was carried out at the transgenic gene-targeting mouse core facility at the University of Utah and we are thankful for their services. We also acknowledge the Cell Imaging core facility and Chris Rodesch at the University of Utah, and the imaging facility at RCB for imaging help, the mouse facility and Penny Noel at the University of Utah for help with mouse colony maintenance, and the genomics core facility at the University of Utah for the RNA sequencing. We thank Jennifer Lawson for important suggestions and help with the mouse work, Eric Bogenschutz and Dr. Santhosh Karanth for suggestions on the RNA-Seq analysis. We are grateful to Prof. Leslie Leinwand for helpful suggestions. We thank Dr. Suchitra Gopinath from THSTI for providing us some antibodies used in this work. We also acknowledge past and present members of GK and SJM labs for valuable suggestions and inputs. Authors acknowledge the support of DBT e-Library Consortium (DeLCON) for providing access to e-resources.

